# Pre-oviposition development of the brown anole (*Anolis sagrei*)

**DOI:** 10.1101/2024.12.15.628549

**Authors:** Antonia Weberling, Natalia A. Shylo, Bonnie K. Kircher, Hannah Wilson, Melainia McClain, Marta Marchini, Katherine Starr, Thomas J. Sanger, Florian Hollfelder, Paul Trainor

**Affiliations:** All Souls College, University of Oxford, Oxford OX1 4AL, United Kingdom; Nuffield Department of Women’s’ and Reproductive Health, University of Oxford, Women’s Centre (Level 3), John Radcliffe Hospital OX3 9DU, United Kingdom; Department of Biochemistry, University of Cambridge, Hopkins Building, Tennis Court Road, Cambridge CB2 1GA, United Kingdom; Stowers Institute for Medical Research, Kansas City, MO, United States; MD Anderson Cancer Center, University of Texas, Texas, United States; Department of Biology, Loyola University Chicago, Chicago, Illinois, USA; Department of Anatomy and Cell Biology, University of Kansas Medical Center, Kansas City, MO, United States

## Abstract

The brown anole, *Anolis sagrei*, has emerged as a representative squamate species during the past several decades. Novel functional tools have been established to manipulate embryogenesis through genome editing or the introduction of small molecule inhibitors, and their effective use requires a thorough understanding of early anole embryogenesis. To enable precise and reproducible staging of anole embryos, we need knowledge of the progression of anole embryogenesis and morphogenesis. While post-oviposition development has been described, the pre-oviposition period remains to be explored. Here, we provide the first staging series of pre-oviposition development for the brown anole. Analysing the follicles and embryos through brightfield imaging, SEM, STEM, histology, and DAPI staining, we define 26 distinct developmental stages. Our dataset reveals that peri-gastrulation morphogenesis up to the initiation of neurulation diverges significantly from chicken, the common representative model of reptile embryogenesis. Furthermore, we followed heart development, neural crest cell migration and central nervous system development through immunofluorescence analyses and provide new comparative insights into the morphogenesis of each of these organ systems over time. Our work establishes the brown anole as a squamate model organism for cross clade evolutionary studies of early embryogenesis.

## Introduction

Early reptile embryogenesis is a severely understudied research area with the most relevant literature dating back to the 19^th^ and beginning of the 20^th^ century.^1^ With the chicken^2,3^ being used as the representative model for avian and non-avian reptile morphogenesis, only limited studies have been carried out on non-avian reptile development.^4^ In 1961, the complete embryogenesis of the lizard *Zootoca (Lacerta) vivipara* was described in 40 stages.^5^ *Agama impalearis*^6^, *Chameleo lateralis*^7^, and *Lacerta agilis*^8^ pre– and post-oviposition development were also documented, but other than these datasets, published staging series focused either on late pre-oviposition or post-oviposition embryogenesis or specific timepoints, that referenced back to chicken development and the *Zootoca* series.^1–4^ This limitation could be due to the inaccessibility of pre-oviposition embryos of many species and the difficulty of staging embryogenesis in the absence of a definitive date of fertilisation as a starting point, as many non-avian reptiles are known to store sperm.^9,10^ Reptiles comprise about 23,000 species and exhibit an incredible range of morphological and ecological adaptations, the basis of which is established during embryogenesis, but remains severely understudied. Increased interest in reptile development and the evolution of developmental mechanisms across distantly related clades^11,12^ requires that we overcome these hurdles and expand the range of research organisms for pre-oviposition development to compare and contrast the mechanisms involved.

Vertebrate development progresses through a series of defined stages – zygote, cleavage, gastrulation, neurulation, and organogenesis, although the details of those stages vary widely among species. For pre– and peri-gastrulation development, we rely often on dorsal brightfield images that provide only limited information about 3D tissue organisation. The chicken embryo undergoes cleavage divisions, forms the area pellucida and then gives rise to the hypoblast resulting in a flat monolayered disc composed of a monolayered epiblast loosely covered with hypoblast cells on its dorsal side overlaying the subgerminal cavity.^2,13^ These developmental steps take roughly 24h and occur inside the mother. The egg is laid as the anterior-posterior axis becomes specified at Eyal-Giladi Kochav (EGK) stage X.^2,14^ During the next 6h, the embryo prepares for gastrulation at EGK stage XIV, also known as Hamburger Hamilton (HH) stage 2. HH1 defines the entire pre-streak development period in this staging system.^2,3,14^ During gastrulation, the primitive streak forms at the posterior border of area pellucida (subgerminal cavity) and extends across about 80% of the embryo before it then regresses again during the next 19h.^15,16^ Neurulation, follows shortly thereafter, during which the head becomes flexed at HH stage 14.^3^ In *Zootoca*, a squamous blastopore is observed at the posterior of the embryonic shield, perpendicular to the anterior-posterior axis of the embryo, and it becomes a large blastoporal canal, forming an elongated tunnel-shaped embryo with the anterior-posterior axis remaining flat until the body rotates at stage 21.^5^

The squamate genus *Anolis* has become a model clade for adaptive radiation since the mid-20^th^ century^17^. With about 400 individual species found across the Caribbean, Central and South America, *Anolis* provides a unique model to study the genomic bases of species diversity.^17,18^ Several *Anolis* species, including *A. sagrei*^19^ and *A. carolinensis*^20^, have had their genomes sequenced^21–23^, opening the door for comparative genomics and functional studies through genome manipulation (e.g., CRISPR/Cas9). As interest in anole development grows, we need a more thorough understanding of the embryogenesis of the brown anole, which has emerged as a model species for the genus. This will allow investigation of the effects of genetic manipulations on embryogenesis as well as enable precise timing for CRISPR/Cas9 injection in developing embryos. While post-oviposition development has been staged^24^, pre-oviposition development remains to be described. Brown and green anoles lay 1 egg every 5-10 days^17,25,26^ throughout the breeding season, from late March to September^24,27^. They also store sperm^28,29^. This prevents an embryo staging system based on timepoint of fertilisation, as has been established for common model organisms, such as mouse and human. Establishing a staging series without knowing the gestational age of the embryos recovered, requires high sample numbers in order to cover all stages by normal distribution across a wild population. The brown anole provides an ideal model organism for the anole group as it is invasive and highly abundant in most of the southern USA, which allows for the collection of large numbers of females.^30,31^

Here, we present the first detailed description of pre-oviposition development of the brown anole. We report 26 distinct stages spanning the first cleavage divisions up to limb-bud initiation, at which point oviposition occurs, and provide insights into establishment of the central nervous system, neural crest cell migration and early cardiac and muscle development. We observed a high degree of divergence from chicken embryogenesis,^2,3^ which is commonly regarded as being representative of avian and non-avian reptiles, as well as from *Zootoca,*^5^ especially at peri-gastrulation stages and the initiation of neurulation. Our staging series not only lays the basis for functional studies of anole embryogenesis but will also enable cross genus and cross-clade comparative studies.

## Results

The brown anole reproductive system has been previously described and consists of two paired ovaries that sit medially to two independent tubes that meet the digestive tract posteriorly to form a single opening called the cloaca. ^28,29^ Each tube of the reproductive tract has three organs including, from anterior to posterior the infundibulum, the glandular uterus, and the nonglandular uterus. The ovaries mature and ovulate follicles in an alternating manner, and the infundibulum is a funnel that receives ovulated ovarian follicles. In the glandular uterus, the eggshell is deposited and fertilised eggs develop. Sperm is stored in the nonglandular uterus and eggs are pushed through this organ and out of the cloaca during laying^28,29^.

To establish a staging series for *A. sagrei* pre-oviposition development, we collected 130 anole females in May 2023 in Southern Florida, USA, and dissected the embryos within 24-96h after capture. Upon dissection, Snout-Vent-Length (SVL, a measurement of body size), number of non-shelled follicles containing yolk, and number of shelled eggs were recorded (Figure 1A). The SVL correlated with the number of shelled eggs and non-shelled follicles containing yolk (Figure 1B). For the following embryonic stages, we first describe our observations based on brightfield images, hematoxylin & eosin staining of paraffin cross sections, electron microscopy imaging, and confocal imaging and then conclude with observed, discernible stages.

**Figure 1:**
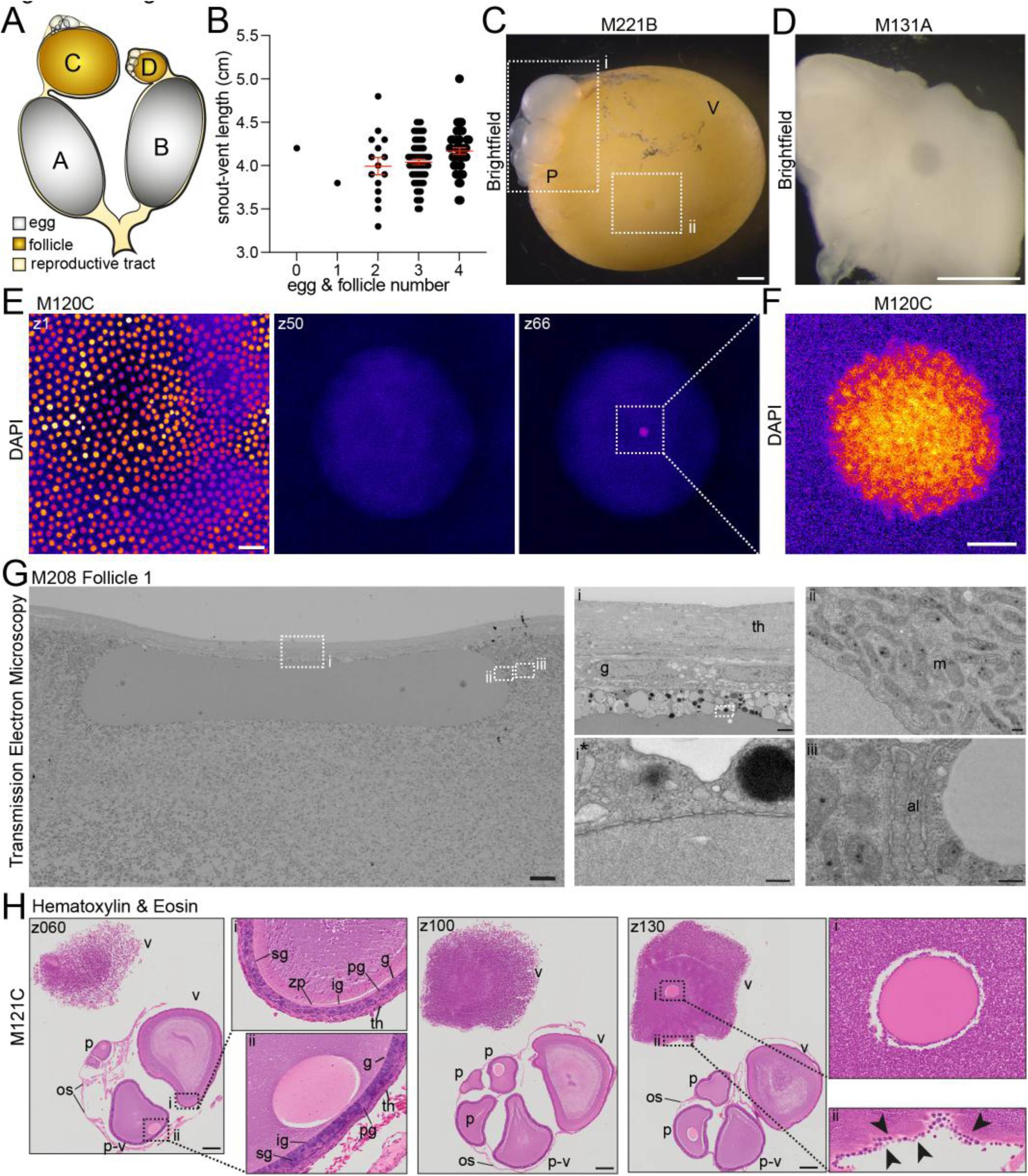
Oogenesis. **A**. Schematic of female reproductive tract. white-grey: eggs, yellow-orange: vitellogenic follicles, light yellow-beige: reproductive tract. Letters A-D illustrate the labelling of embryos and follicles depicted from a single female with A labelling the largest egg/follicle, B the second largest etc. **B.** Analysis of female snout to vent length (SVL) and number of eggs & yellow yolk-filled vitellogenic follicles. Scatter plot, mean±SEM in red. each dot represents one female. **C.** Brightfield image of a vitellogenic yolk-filled follicle (V) M221B with ovary and previtellogenic follicles (P) attached. Dotted rectangles outline (i) previtellogenic follicles, (ii) position of the germinal vesicle. Scale bar: 1mm. **D.** Brightfield image of dissected nucleus from M131A. Scale bar: 1mm. **E.** DAPI staining of M120C. left z1, middle z50, right z66. Scale bar 40um. **F**. DAPI staining in the centre of nucleus. Scale bar 5um. **G.** Transmission electron microscopy images of nucleus of M208 follicle 1. Left: full nucleus. Right: zoom-ins of regions of interest indicated by i/ii & */**.i multiple cell layers cover the follicle. ii the nucleus is surrounded by mitochondria. * nuclear pores. ** nuclear pore complexes. scale bars: left = 20um, i = 2um, ii/*/** = 200nm.**H.** Cross sections of M121C. Hematoxylin & Eosin staining. Sections z060, z100, z130 are shown, full z-stack in Figure S1. z060 arrows highlight membrane enclosing ovary. Previtellogenic follicle (P), vitellogenic follicle (V), pre-to vitellogenic follicle (P-V). Boxes (i) zoom in of cell layers surrounding individual follicles. Arrows highlight multilayer of cells. (ii) Scale bar:250um

## Oogenesis

We initially focussed on the ovaries and the large yolk-filled vitellogenic follicles that had not yet ovulated. Each of the two anole ovaries contains follicles with increasing volume. Typically, we observed 1-2 vitellogenic follicles, the late stage of which are significantly larger than early vitellogenic follicles and the previtellogenic follicles (Figure 1Ci, Figure S1A). The late yolk-filled vitellogenic follicle is bright yellow and exhibits a round area of much lighter colour, the germinal disc.^32^ In the middle of the germinal disc at the position of the nucleus or germinal vesicle,^32^ a round, dark shadow can be distinguished (Figure 1Cii). This area could be dissected from the follicle and contained a defined spherical structure (Figure 1D). To confirm that this structure is indeed the nucleus of the oocyte, we DAPI stained the dissected structures (Figure 1E) and performed 3D imaging. The late vitellogenic follicles are surrounded by a single cell layer of follicular epithelium (Figure 1E, z1), and we observed a DAPI positive area in the middle of the germinal vesicle (Figure 1E, z66). When taking a high-resolution image of the DAPI positive area, it became clear that it was not a solid structure but rather exhibited varying levels of intensity reflecting different concentrations of DNA (Figure 1F) and thus could be the nucleolus. We performed transmission electron microscopy on the nucleus of an early vitellogenic follicle to determine the organisation of subcellular structures around the nucleus. The pre-processing resulted in the sphere becoming flat and oval shaped, the imaging of which revealed that the entire nucleus was devoid of organelles and other intracellular structures (Figure 1G). Furthermore, it is covered by several cell layers, of outer theca and inner granulosa cells, and surrounded by lipid and yolk droplets (Figure 1Gi). When analysing the membrane surrounding the nucleus under higher magnification, we found that it consists of a double layer and includes a number of pores (Figure 1Gi*). Double membranes with pore complexes dispersed throughout are a hallmark for the lamina that surround nuclei. We next focussed on the structures adjacent to the nucleus and found a high concentration of mitochondria and nuclear pore complexes as well as annulate lamellae (Figure 1Gi,i**). Taken together, these results confirm that the entire sphere of about 150μm diameter is the germinal vesicle.^33^

To better define the anatomy of the follicles in more detail^34^, we sectioned an ovary and stained it with hematoxylin and eosin (H&E) (Figure 1H, Figure S1B). The largest vitellogenic follicle broke apart during processing, but the previtellogenic follicles and the early vitellogenic follicles remained intact and are surrounded by a thin membrane, the ovarian stroma (Figure 1H z060, arrows). The individual follicles are surrounded by the zona pellucida and covered by a thick multicellular envelope that consists of two distinct cell types^35^, confirming our observations from STEM (Figure 1Gi, Figure 1H z060i). The granulosa cells directly overlie the zona pellucida and form the inner layers visible as cuboidal cells^32^. Three different types of granulosa cells can be distinguished; small, intermediate, and pyriform (Figure 1H z060 i/ii).^36^ The outer layer is composed of longitudinal cells, the theca cells^37^ (Figure 1H z060i/ii). Each follicle contains a spherical nucleus located right beneath the multicellular envelope, which does not exhibit any internal features (Figure 1H z060ii, z100, z130i) supporting our findings from STEM. Although the largest vitellogenic follicle was damaged during the processing, we could observe parts of the follicular epithelium (Figure H z130i, arrows) as well as the nucleus.

## Cleavage Divisions

We next analysed eggs dissected from the glandular uterus.^29^ Each of them was enveloped by a shell, which ranged from a very thin, skin-like layer to a fully mature white eggshell. Following fertilisation, the anole embryo undergoes meroblastic cleavage divisions, with blastomeres dividing on top of the large yolk. The first two cleavage furrows appear in the middle of the embryonic disc, partitioning the embryo into 4 blastomeres (Figure 2A M239B, arrows). Then, the first large blastomeres appear in the middle with the furrows extending from the blastomeres towards the outer border of the embryonic disc (Figure 2A M151B, arrows). The further the cleavage divisions progress, the smaller the blastomeres become until the entire embryonic disc is covered with small blastomeres (Figure 2A M129A). To investigate the process of the initial cleavage divisions in more detail, we performed scanning electron microscopy (SEM) (Figure 2B left). When the first 2 furrows are established, the next 4 furrows form to mediate the subsequent cleavages. These furrows initiate from two sides. Close to the middle of the embryo, a ridge is established flanked by two indentations (Figure 2Bi, i*, i**). Towards the outside and in a comparable distance from the midpoint as the established furrows, an invagination forms (Figure 2Bii). Notably, we observed that the ridge and the inside of the established furrow exhibit similar surface structures (Figure 2Bi**,iii, iii***). The mechanism by which the two parts of the cleavage furrow fuse remains to be described.

**Figure 2:**
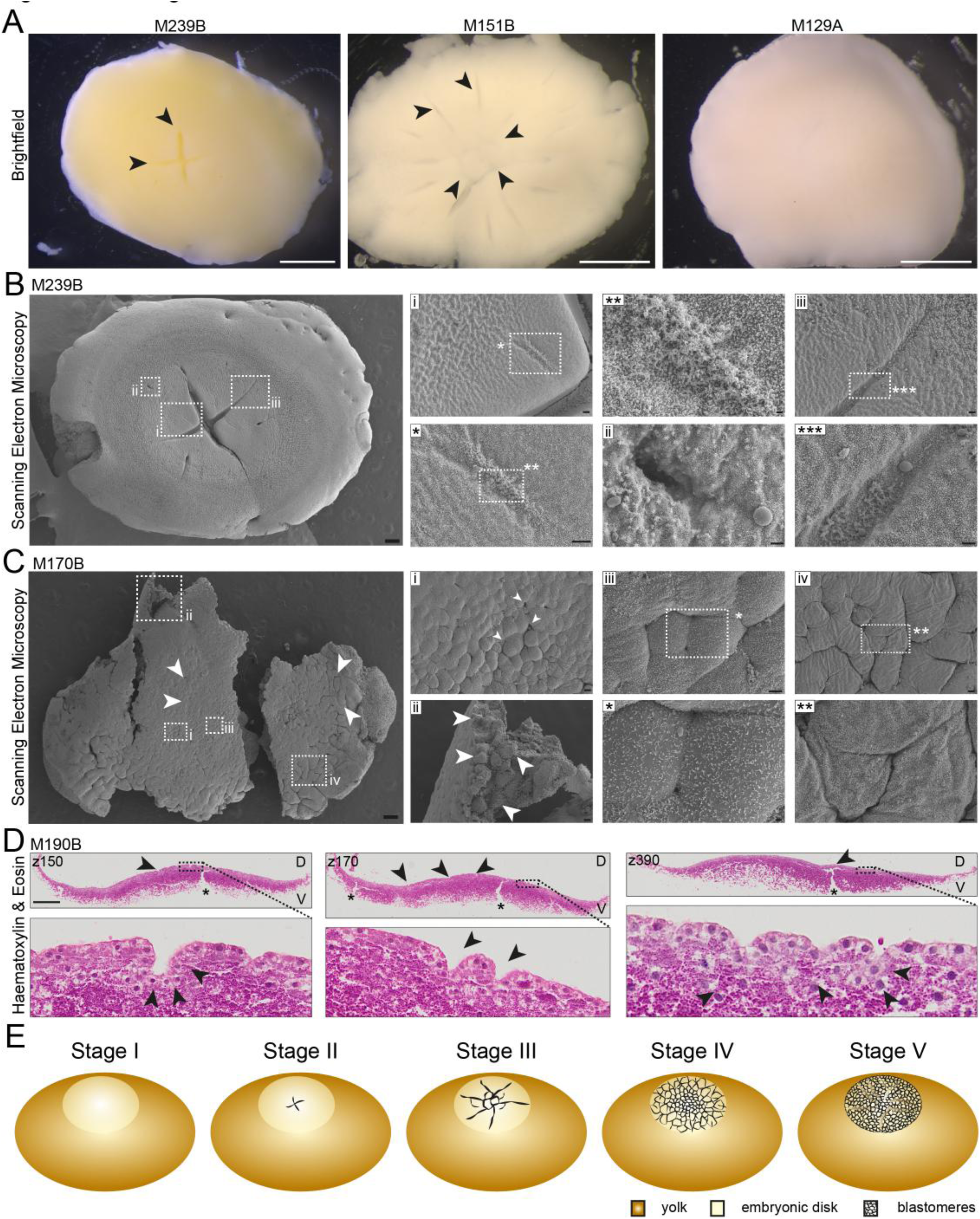
Cleavage Divisions. **A.** Brightfield images of embryos M239B, M151B, M129B (left to right). arrows highlight cleavage furrows (M239B) and furrows and initial blastomeres (M151B). scale bars 1mm. **B.** Scanning electron microscopy imaging of early cleavage stage embryo M239B exhibiting 2 mature furrows. Red boxes highlight zoom in regions labelled with i/ii/iii/*/**/***. (i/*/**) ridge at the central cleavage furrow initiation. (ii) shows the outer cleavage furrow initiation. (iii/***) mature cleavage furrow focussing on the surface structure of the furrow surface (***). scale bars: left = 200um, i/iii/* = 20um, ii/*** = 10um, ** = 2um. **C.** Scanning electron microscopy imaging of late cleavage stage embryo M170B. centre formed of small blastomeres, the outer layers contain larger cells with cleavage furrows and incomplete cellularisation visible at the outer border of the embryo (arrows). boxes depict zoom in regions i-iv/*/**. (i/iii) central blastomeres of different sizes, arrows highlight points of possible cell internalisation, (iii box *) incomplete cell cleavage in embryo centre. (ii) multiple cell layers beneath the surface. arrows highlight inner and outer cells. (iv/**) cleavage furrow in outer region of embryo. scale bars: left 200um, i/ii/iv = 20um, iii/** = 10um, * = 2um. **D.** Hematoxylin and Eosin staining of paraffin cross sections of cleavage stage embryo M190B, z150, z170, z390 shown. D marks the dorsal, V the ventral side. Full z-stack in Figure S2. Boxes outline zoom in regions in bottom row. Asterisk: regions of yolk breakage. Arrows top row: cleavage furrow. Arrows bottom rom cleavage furrows composed of multiple cells (z150, z170), internalised cells (z390). Scale bar: 400um. Full z-stack in Figure S3. **E.** schematic of embryo stages I-V.

We next analysed a later cleavage stage embryo by SEM. Here, the entire embryonic disc is covered by blastomeres (Figure 2C). In the middle, the blastomeres are smaller than those at the edge of the embryonic disc (Figure 2C, arrows). At the edge of the embryonic disc, cleavage furrows remain, indicating that the cleavages have not yet completed. Instead of forming one flat layer, the cells in the middle appear to overlay each other at several points, where cell internalisation could have taken place (Figure 2Ci, arrows). The cross-break of the embryo reveals several layers of cells at this stage (Figure 2Cii, arrows). In the middle of the embryo, the blastomeres while overall significantly smaller than at the edge of the embryonic disc, exhibit a high degree of size variation, likely due to ongoing cell cleavage (Figure 2Ci), as evidenced by cleavage furrows (Figure 2Ciii, iii*). A similar phenotype could be observed at the outer region of the embryo (Figure 2Civ, iv**). To gain further information on the morphology of cleavage stage embryos and whether the multilayer found in Figure 2Cii was truly formed by multiple cells, we sectioned cleavage stage embryo M190B and stained with H&E (Figure 2D, Figure S2A/B). Several large furrows were observed (Figure 2D z150, z170, z390, arrows top row). At this stage, the embryo is not a cohesive structure but rather a loose flat structure on top of the yolk which can only be dissected by retaining a layer of yolk beneath. Yolk breakage points (Figure 2D z150, z170, z390, asterisks top row) are not connected to the cleavage furrows. The furrows are composed of multiple cells arranged to form an indentation (Figure 2D z150, z170, z390, bottom row). We observe multiple cells beneath the top cell layer (Figure 2D z390, bottom, arrows), though whether these localise there through asymmetric cleavage or internalisation remains to be elucidated.

**Stage I:** zygote

**Stage II:** the first cleavage furrows appear in the middle of the embryo but does not extend over the entire embryonic disk.

**Stage III:** the first defined large blastomeres form in the middle of the embryo with the furrows extending further towards the edges of the embryonic disk.

**Stage IV:** blastomeres can be found all over the embryonic disc with small blastomeres in the middle and larger ones towards the outside with cleavage furrows still extending at the edges of the embryonic disk. Some blastomeres are found in a second or third layer.

**Stage V**: the embryo is comprised of tiny, bead-like cells with multiple cells forming furrows and extending into the yolk.

At none of these stages can the embryo be recovered without a layer of yolk underneath. (Figure 2E, Table S1)

## Peri-gastrulation development

Upon conclusion of blastomere cleavage, the embryo is not yet a cohesive structure. This changes as development progresses and the embryo becomes one coherent sheet-like structure encompassing the entire embryonic shield with a slight thickening in the middle (Figure 3A M126B). We could not observe a subgerminal cavity as is found in chicken^13^, instead, the multilayered embryo lies directly on top of the yolk. Shortly thereafter, the anole embryo adopts a dome-like configuration with a shallow curvature at first (Figure 3A M184B), which then becomes highly convex while remaining radially symmetric (Figure 3A M142B). This symmetry breaks during anterior-posterior patterning by a slight thickening of the posterior side (Figure 3A M144B) which becomes more pronounced during gastrulation (Figure 3A M123A). To determine whether this structure was fully composed of cells, we imaged the ventral side and observed that once yolk is cleared away, the embryo forms a hollow dome (Figure 3A, bottom row) which directly overlays the yolk and exhibits a higher curvature than the overall curvature of the egg. To gather more information on the cellular composition of the embryo at these stages, we performed H&E staining on cross sections (Figures 3B, S3A/B). This confirmed our observations of the overall curvature of the embryo. Progressing through the sectioned embryo, the morphology is initially flat, and then the embryo exhibits a distinct curvature that decreases at its posterior end. We could observe 2-3 cell layers across the embryo in general (arrows) and a pronounced multilayered section of up to 8 cell layers (Figures 3B z105, S3B z090-132).

**Figure 3:**
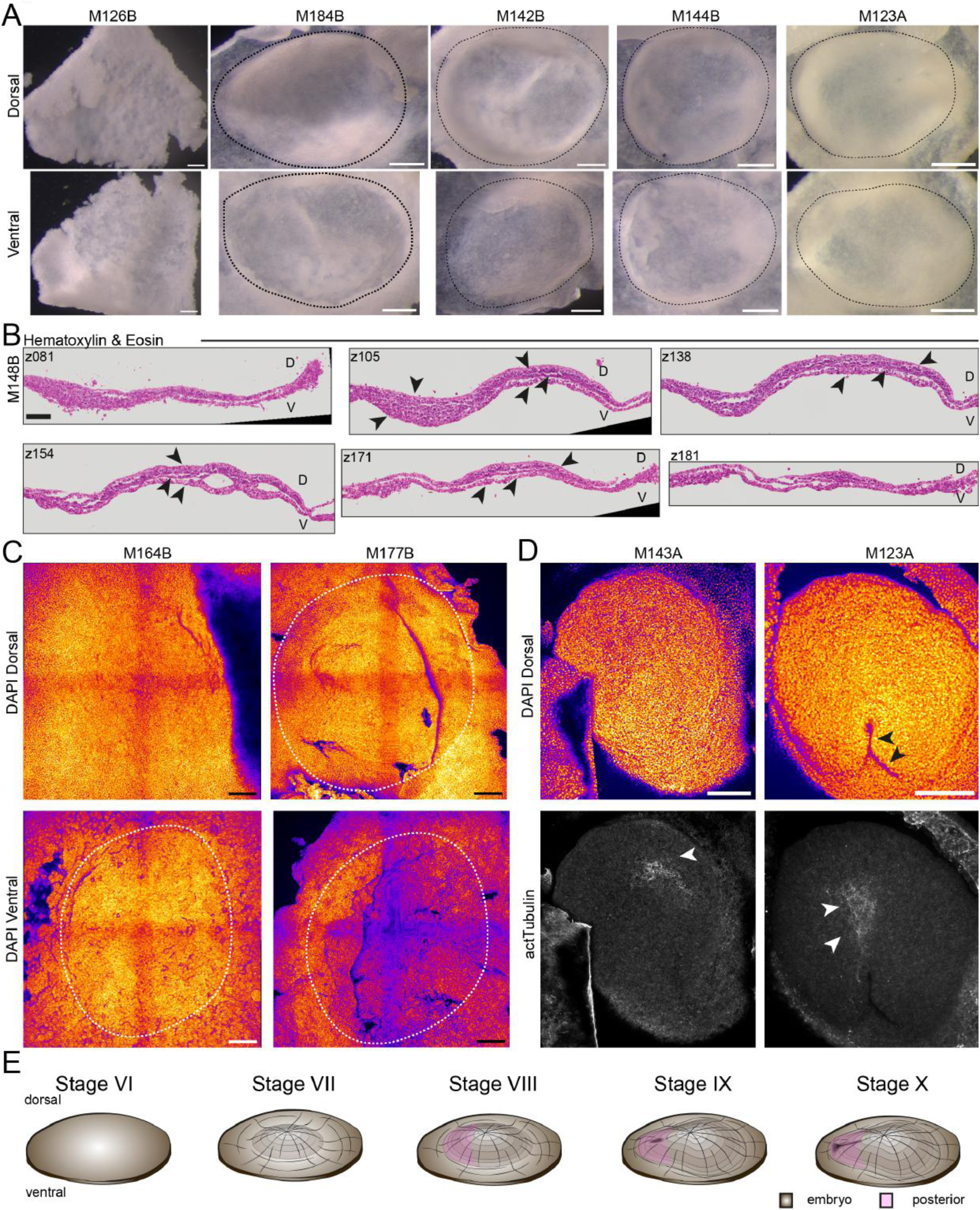
Perigastrulation. **A**. Brightfield images of peri-gastrulation stage embryos M126B, M184B, M142B, M144B, M123A. Embryos exhibit increasing curvature forming dome-shaped structures and are outlined with dashed lines. top row dorsal, bottom row ventral view. Scale bars 250um. **B.** Hematoxylin and Eosin staining of paraffin cross sections of peri-gastrulation stage embryo M148B. sections z081, z105, z138, z154, z171, z181 shown, Full z-stack in Figure S3B. Embryo consists of multiple cell layers (arrows). scale bar: 100um. C. Maximum Intensity projection of DAPI stained embryos 164B, M177B. top row dorsal, bottom row ventral view. Dashed outline around the embryo. scale bars: 250um. **D.** Maximum Intensity projections of dorsal view of DAPI and actTubulin stained embryos M143A, M123A. top row arrows depict blastopore. Bottom row arrows highlight expression of actTubulin. Scale bars:250um. Fire staining corresponds to signal intensity with yellow as high and purple as low signal. **E.** schematic of embryonic stages VI-X.

We then performed pseudo-SEM^38,39^ on embryos of consecutive developmental stages. We confirmed the formation of a flat sheet (Figure 3C M164B dorsal) with a thickening in the middle region on the ventral side (Figure 3C M164B ventral, dashed outline. Then the dome-like curvature initiates (Figure 3C M177B, dashed outline). Following the formation of a highly curved dome (Figure 3D M143A), the embryo starts to exhibit anterior-posterior asymmetry. This is initially visible through asymmetric distribution of acetylated Tubulin, which labels prospective anterior neural tissue (Figure 3D bottom row), as the embryo becomes oval shaped. This asymmetry becomes more pronounced with the posterior end developing with a cone-like tip (Figure S3C M141A) which exhibits a slight indentation (arrow), consistent with being the blastopore. The blastopore then develops into a clear groove that extends towards the anterior end of the embryo (Figure 3D M123A, top arrows). Acetylated Tubulin staining remains anterior of this structure and is now found more towards the middle of the embryo (Figure 3D M123A, bottom) marking the prospective neural groove. Taken together, the anole embryo forms a highly curved, hollow, dome-like structure upon gastrulation with its convex side dorsally (Figure 3E).

**Stage VI:** The embryo becomes one coherent flat sheet.

**Stage VII:** The embryo is a dome shaped with a flat, symmetrical curvature.

**Stage VIII:** Curvature increases while the embryo remains overall symmetrical but starts to exhibit slight anterior-posterior asymmetry.

**Stage IX:** Anterior-posterior asymmetry is established with a posterior thickening and appearance of the blastopore.

**Stage X:** The blastopore then develops into a groove that grows towards the anterior (Figure 3E).

## Initiation of Neurulation

Following gastrulation, the anole initiates neurulation. At this stage, the neural groove extends from anterior to posterior, while the neural folds do not appear to be elevated yet (Figures 4A/S4A M149B). The head field remains flat and covered by the amnion. The embryo retains its dorsally highly convex shape acquired during gastrulation with the middle of the developing embryo being the most elevated region. Then, the neural folds elevate to initiate neural tube closure in the middle of the embryo, before extending anteriorly and posteriorly with the head field remaining flat (Figures 4A/S4A M242A). Next, the head folds become elevated, and the embryo loses its anterior to posterior curvature, becoming flat while the amnion now extends from the anterior up to the middle of the embryo (Figures 4A/S4A, M250B).

**Figure 4:**
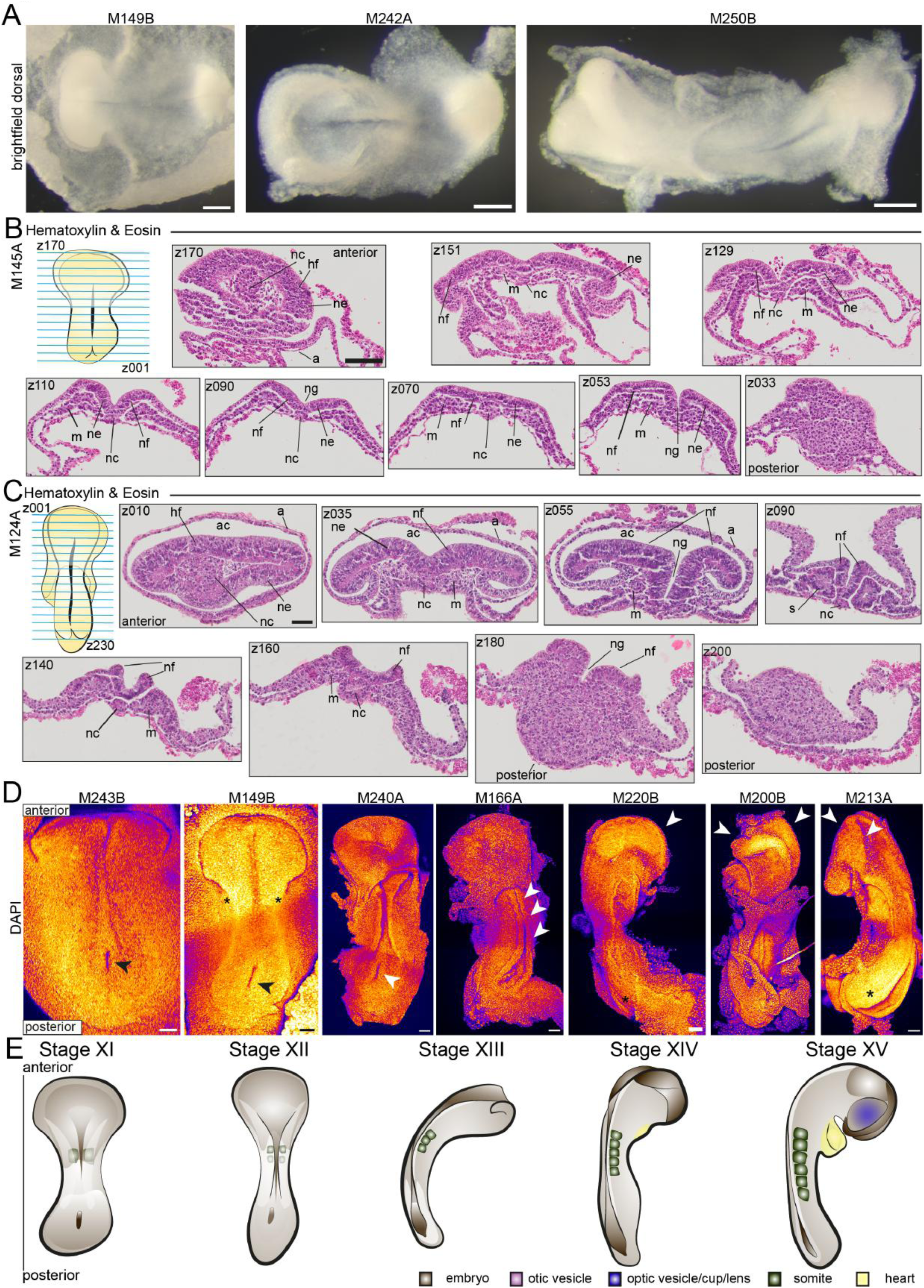
Initiation of Neurulation. **A**. Brightfield images of early neurulating embryos M149B, M242B, M215B. Dorsal views. ventral views in Figure S4A. Scale bar 200um. **B.** Hematoxylin and Eosin staining of paraffin cross sections of embryo M145A. top row left schematic drawing of embryo, blue lines symbolise cross sections z170-z001 indicate the direction of sectioning. From anterior to posterior sections z170, z151, z129, z110, z090, z070, z053, z033 are shown. Full z-stack in Figure S5B. annotations: a= amnion, hf = head field, m = mesoderm, nc = notochord, ne = neuroepithelium, nf = neural fold, ng = neural groove. Scale bars 100um. **C.** Hematoxylin and Eosin staining of paraffin cross sections of embryo M124A. top row left schematic of sectioned embryo with blue lines depicting cross sections and z001-z230 showing direction of sectioning. From anterior to posterior sections z010, z035, z055, z090, z140, z160, z180, z200 are shown. Full z-stack in Figure S6B. Annotations as in (B) plus: ac=amniotic cavity, s = somite. Scale bars 100um. **D.** Maximum Intensity projections of DAPI stained embryos M243B, M149B, M240A, M166A, M220B, M200B, M213A. dorsal view. Anterior top, posterior bottom. M243B, M149B, M240A arrow: blastopore. M149B asterisk: position of first developing somites. M166A arrows: neural tube closure in middle of embryo. M220B, M200B, M213A arrows: head folds. M220B, M213A asterisk: posterior neuropore. Scale bars 100um. Fire staining corresponds to signal intensity with yellow as high and purple as low signal. **E.** Schematic of embryonic stages XI-XV.

We stained cross sections of embryos at consecutive stages of early neurulation with H&E (Figures 4B/S5A/B M145A, 4C/S6A/B 124A, S7A/B M134B). In M145A, the notochord and flat head field are clearly visible (z170). The median hinge point is well formed in the anterior neural plate, with neural folds becoming elevated (z110-090), but more caudally the neural plate remains flat (z070). The neural groove is visible in both the anterior and the posterior of the embryo (z053) (Figures 4B/S5B). Following this stage, the neural folds elevate along the entire embryo except for the anterior of the head field (z010) in M124A (Figures 4C, S6A/B). The neural folds appear to close like flat sheets in the middle of the embryo (z090), while they elevate in a rounder manner towards the posterior as the neural tube forms (z140, z160). Somitogenesis has also commenced by this stage (z090). Then, the head folds (Figure S7A/B z004-029) and the neural tube closes in the middle of the embryo (z054-z170) while the anterior and posterior neuropores remain open.

To gain further insights into the initiation of neurulation in anoles, we performed pseudo-SEM staining of embryos at consecutive stages of development (Figure 4D). The blastopore (neurenteric canal) that forms during the late stages of gastrulation (Figure 3D M141A) is present until the neural tube closes in the middle of the embryo (Figure 4D M243B, M149B, M240A, arrow). We observed initial fusion of the neural tube in the middle of the embryo, which then extends rostrally and caudally. We also observed the initiation of somitogenesis (somite formation) in embryos that exhibited elevation of the neural folds (Figure 4D, M149B star, Figure S8A/B). Following the initiation of neural tube closure (Figure 4D M166A, arrows), the head folds begin elevating with the right and left sides turned laterally (Figure 4D M220B, M200B, M213A, arrows). Similarly, elevation of the neural folds extends posteriorly with the posterior neuropore remaining open (Figure 4D, M20B, M213A asterisk). The embryo initiates embryonic turning to the right after becoming flat during neural tube closure (Figure 4D M240A, M166A (flat), M213A (turned)), which positions its left side against the yolk, as has been reported for sauropsidae.^3,5,24,40,41^

**Stage XI:** The embryo is highly curved and exhibits a neural groove with neural folds that are slightly elevated in the middle of the embryo but not bent towards each other yet, and flat in the anterior and posterior of the embryo. The embryo has up to one somite.

**Stage XII:** The neural folds are elevated along the length of the embryo and bent towards each other except for the anterior and posterior, which remain flat. The amnion extends over the head field and the embryo has up to 2 somites. The embryo has lost its overall curvature.

**Stage XIII:** The neural tube closes, beginning in the middle of the embryo, while the anterior and posterior neuropores remain wide open. The head folds are either still flat and open or have initiated folding, and the embryo has 3 somites.

**Stage XIV:** The embryo initiates bending from the head folds to the tail bud and has 4-5 somites. The neural tube is closed in the middle, while the anterior and posterior neuropores remain wide open. The tail bud appears and is angled slightly inward to the midline. The head starts to tilt while the head folds start to bend towards each other. The nascent heart is visible under the head folds.

**Stage XV:** The embryo has 6 somites and the optic vesicles manifest as slight swellings. The anterior and posterior neuropores remain open and the embryo is curved overall again (Figure 4E, Table S1). The heart is detectable as small tube directly under the head folds.

## Completion of Neurulation

Following the initiation of neural tube closure, optic (arrows) and otic (asterisk) vesicle formation occurs together with further development of the heart (outlines) (Figure 5A) and continued somitogenesis. First, the posterior neuropore closes, while the anterior remains wide open (Figure 5A M170A). The optic vesicles are visible as slight swellings and the developing heart tube forms right beneath the head folds prior to embryo turning. The optic vesicles then gradually become more pronounced, and the heart enlarges to become a clearly distinguishable organ (Figure 5A M138A). As the optic cup invaginates, the heart continues to increase in size, and the somites become more easily distinguishable (Figure 5A M235B), and extends more posteriorly as the tail grows and curls. Simultaneously, the first pharyngal arches develop (Figure 5A M138A, M235B).

**Figure 5:**
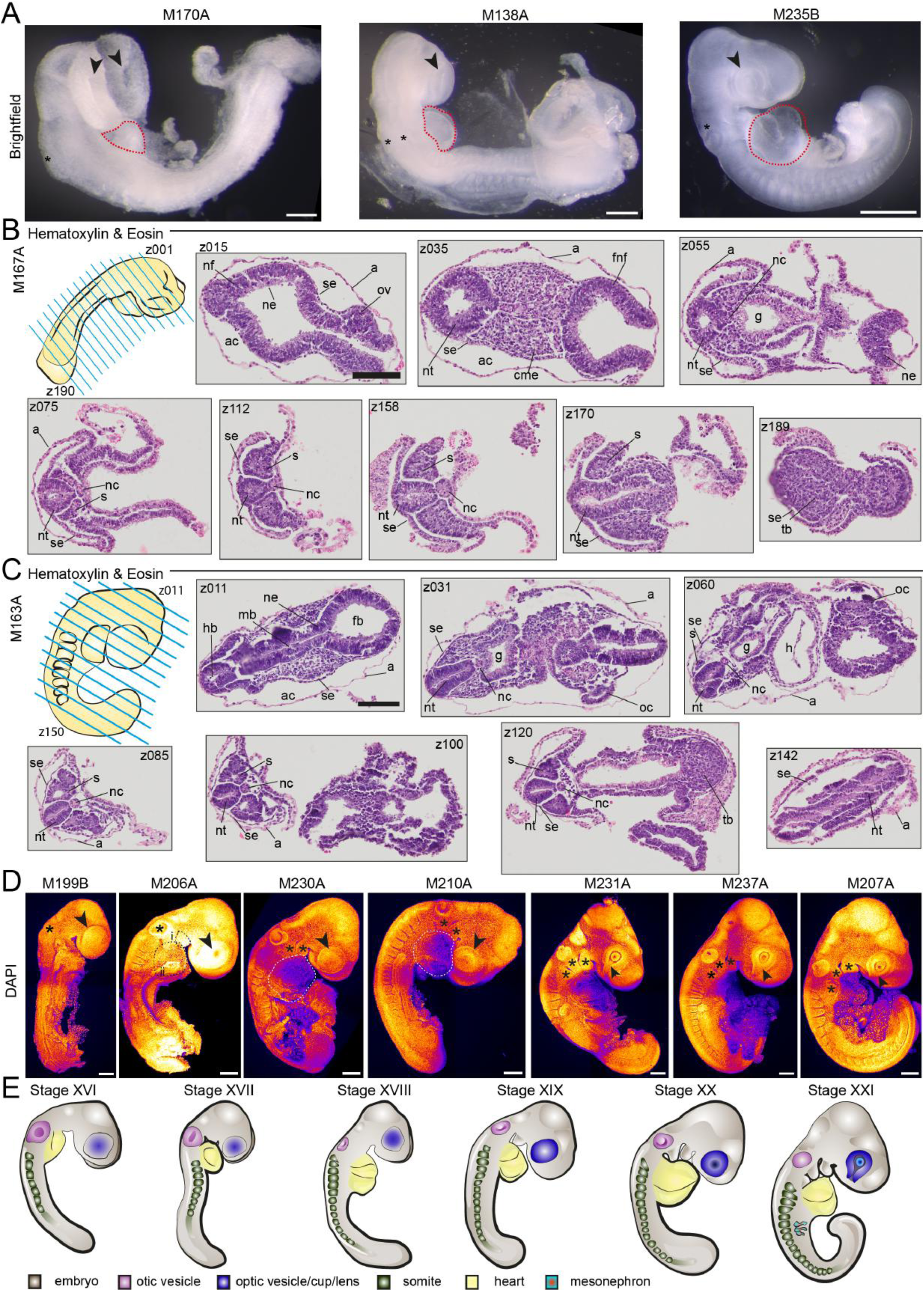
Neurulation. **A**. Brightfield images of M170A, M138A, M235B. Scale bars: M170A/M138A 200um, M235B 500um. **B/C**. Hematoxylin and Eosin staining of paraffin cross section of embryos M167A (B), M163A (C). top rows left schematic drawing of embryo with blue lines indicating cross sections and z001 – z190 (B)/z150 (C) indicate direction of sectioning. (B) sections z015, z035, z055, z075, z112, z158, z170, z189 shown, full z-stack in Figure S9B. Scale bar: 100um. (C) sections z011, z031, z060, z085, z100, z120, z142 shown, full z-stack in Figure S11B. Scale bar: 100um. (B/C) annotations: a= amnion, ac= amnionic cavity, fnf=forebrain neural fold, g = gut, h = heart, nc= notochord, ne = neural epithelium, nf= neural fold, nt= neural tube, ov= optic vesicle swelling, pm = paraxial mesoderm, s= somite, se= surface ectoderm, tb= tail bud. **D.** Maximum Intensity Projections of DAPI staining of embryos M199B, M206A, M230A, M210A, M231A, M237A, M207A. M199B arrow: optic vesicle swelling. M199B, M206A star: otic vesicle. M206A i: pharyngal arch, ii: heart. M230A, M210A asterisk: pharyngal arches, arrows: optic cup, outline: heart. M231A, M237A, M207A asterisk: pharyngal arches, arrow: optic fissure. Scale bars: 200um. Fire staining corresponds to signal intensity with yellow as high and purple as low signal. **E.** Schematic of embryonic stages XVI-XXI.

We then performed H&E staining on cross sections of 3 embryos of consecutive stages to follow the progression of neurulation and initial organ formation in more detail (Figures 5B,S9A/B M167A, S11A/B M152A, 5C,S11A/B M163A). In M167A, the amnion encloses the head folds, which are not yet fused. The optic vesicles are visible as swellings and the neural tube exhibits a rhomboid shape anteriorly with a rhomboid lumen (Figure 5B z035, S9 z035-045), both of which become triangular (Figure 5B z055, S9 z055-065). Moving further from the anterior to the posterior of the embryo, this triangular shape then becomes slightly more spherical while the lumen adopts an elliptical shape (z070-149). At the same time, both notochord and somites are visible. In the following more posterior sections, the neural tube is triangular again, elongated along the dorso-ventral axis, which could also account for the slit-like lumen (z158). The notochord and somites are pronounced. Then due to the overall curvature of the embryo, the neural tube appears as a canal and the last somites are visible but no longer completely detached from the neuroectoderm (z170). This is indicative of the location of neuromesodermal progenitors and the presomitic mesoderm. In the most posterior sections, the tail bud becomes visible (z189). Analysing the next embryo (M152A), we observed the nascent heart (Figure S10B z040-060). Moving further posteriorly, the gut is evident (z020-055) and the embryo is more elongated than at the earlier timepoint (M167A), as is evidenced by an increased number of sections containing the neural tube, notochord and somites only (z070-145). At the latest timepoint studied (M163A), the forebrain, midbrain and hindbrain become distinguishable (Figure 5C, z011, S11A/B M163A), while the optic cup continues to invaginate in concert with lens formation (z031, z060). The gut is now more pronounced than in M152 (z020-060) and the overall body shape is more curved (Figure S11A).

Following this analysis, we carried out pseudo-SEM staining of embryos at consecutive developmental timepoints (Figure 5D). M199B exhibits an open anterior neuropore, while the posterior neuropore is already closed. Anteriorly, the optic vesicles are visible as pronounced swellings (arrow) and the otic vesicle can be discriminated as a slight indentation (asterisk). The somites are clearly apparent at the posterior end, and the tail bud is visible. In M206A, the optic cup has initiated invagination (arrow) and the first pharyngal arch (dotted outline i) as well as the heart have begun to form (dotted outline ii) (Figure 5D M206A). The otic vesicle is clearly visible (asterisk) and the embryo has started to bend from anterior to posterior. The embryo M230A/M210A continues to elongate, as the heart grows significantly in size (white dotted outline), while the second pharyngal arch forms (asterisks) and the optic vesicle (arrow) further invaginates. The lens forms with the optic cup exhibiting a wide optic fissure (arrow) as the next (third) pharyngal arch emerges (asterisks) (M231A, M237A, M207A). Simultaneously, the tail continues to elongate.

**Stage XVI:** The posterior neuropore has closed, while the anterior neuropore remains open, and the optic vesicles are visible as clear swellings. The nascent heart is small and visible, but often withdrawn from direct observation through the amnion. Up to 7 somites.

**Stage XVII:** The 1^st^ pharyngal arch is apparent, and the optic vesicles have become even more pronounced. The otic vesicles have begun invaginating. The embryo is straight overall, but the head is tilted towards the ventral side. 8-9 somites.

**Stage XVIII:** The 1^st^ pharyngal arch is fully formed, the body starts to flex, and the heart is now clearly distinguishable as a small sac. Up to 13 somites.

**Stage XIX:** The anterior neuropore is closed, and olfactory placode formation has begun. Up to 14 somites.

**Stage XX:** The 2^nd^ pharyngal arch has formed, the optic vesicles have begun invaginating, and the lens develops with a wide optic fissure. The tail bud is clearly visible. About 15 somites.

**Stage XXI:** The 3^rd^ pharyngal arch becomes apparent, and the lens forms with an optic fissure. The olfactory placode is clearly visible, the tail starts to curl, and the mesonephri emerge. Up to 19 somites. (Figure 5E, Table S1).

## Organogenesis

Organogenesis begins prior to oviposition, and we observed a significant increase in the size of the head compared to the rest of the body, together with increased growth of the midbrain (red outline). Simultaneously, the next posterior pharyngal arches form (asterisks), the limb buds become visible (blue highlight) and the tail elongates. The latest stage to be dissected exhibited initiation of eye pigmentation (Figure 6A M198A, arrow). When analysing these final stages, we observed a narrowing of the optic fissure (Figure 6B arrows, Figure S12 A-C).The olfactory placode becomes visible as a distinct indentation adjacent to the eye (Figure 6B, M192B asterisk) and the somites extend throughout almost the entire tail. We then carried out H&E analysis on two sectioned embryos (Figures 6C, S13A/B M211A, Figures 6D, S14A/B M225A). In M211A (Figure 6C), we observed the lumens of the prospective fore-mid– and hindbrain formed by the neuroectoderm (Figure 6C, z011, S13, z007-033). This is overlaid by cephalic mesenchyme, which in turn is enveloped by surface ectoderm and on top the amnion (z011/z041). The rostral extremity of the notochord appears in the mesenchymal tissue, directly adjacent to the rostral extent of the foregut which is flanked by the dorsal aorta and the first branchial arch artery (asterisks) (z041). At this time, the optic vesicles are visible (z068-087) and the heart is located directly beneath the head (h) (z068-105). The neural tube is visible as an elliptic structure with a slit-like lumen, flanked by somites and adjacent to the notochord, which lies in close proximity to the hindgut diverticulum (z076-109). The nephric tract starts developing (z121-185) and the elongating tail bud (tb) is clearly visible (z414-449) (Figure S13B).

**Figure 6:**
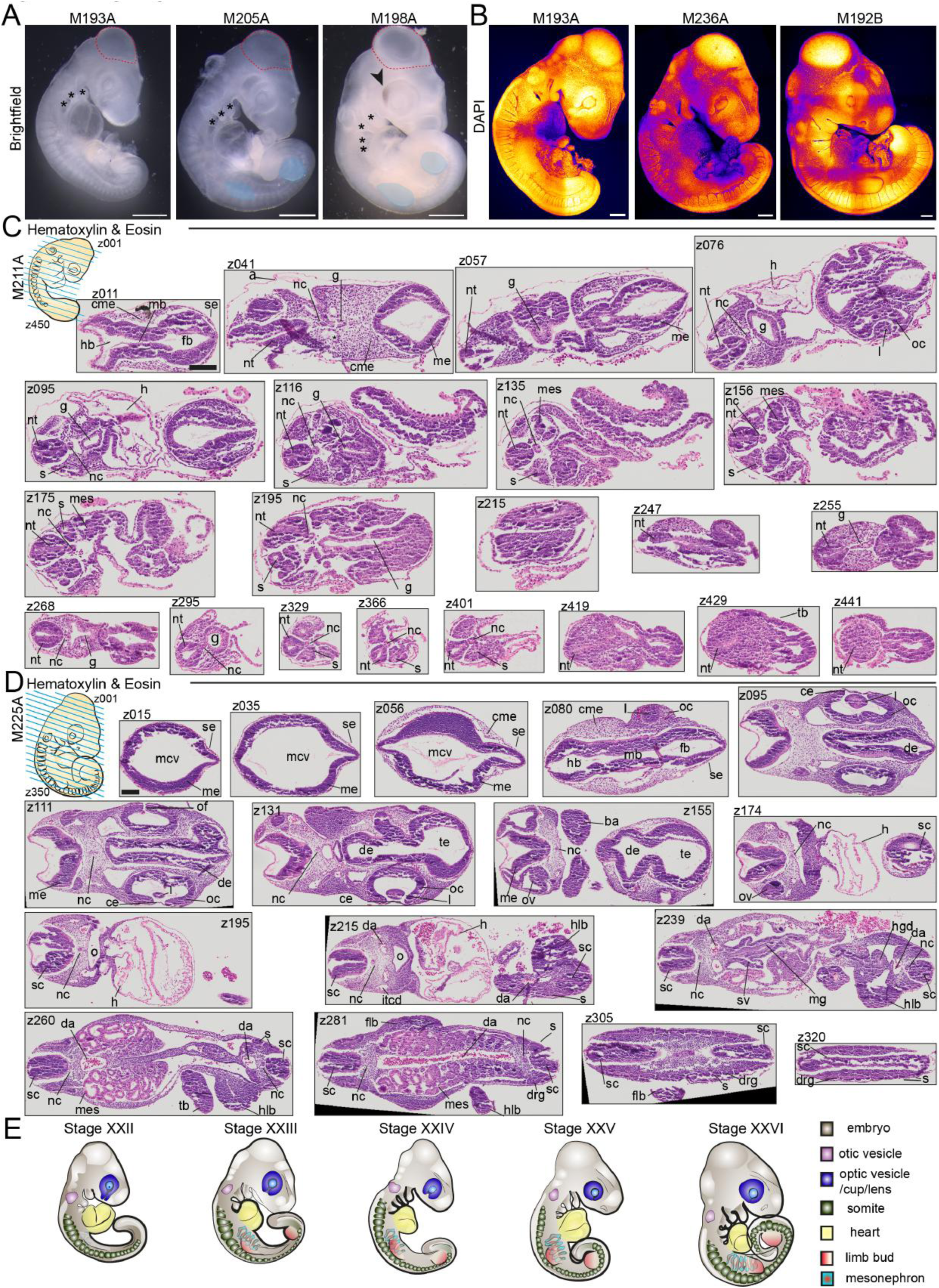
Organogenesis. **A.** Brightfield images of embryos M193A, M205A, M198A. asterisk: pharyngal arches; arrow: pigmentation of eye red outline: midbrain vesicle; blue highlight: limb buds. scale bars 500um. **B.** maximum intensity projections of DAPI staining of embryos M193A, M236A, M192B. arrows: optic fissure; star: olfactory placode; Scale bars 200um. Fire staining corresponds to signal intensity with yellow as high and purple as low signal. **C/D.** Hematoxylin and Eosin staining of paraffin cross sections of embryos M211A (C), M225A (D). left top row schematic drawing of embryo with blue line indicating cross sections and z001-z450 (C)/z350 (D) indicating direction of sectioning. (C) sections z011, z041, z051, z076, z095, z116, z135, z156, z175, z195, z215, z247, z255, z268, z298, z319, z366, z401, z429, z441 shown, full z-stack in Figure S13B. scale bar 100um (D) sections z015, z035, z056, z080, z095, z111, z131, z155, z174, z195, z215, z239, z260, z281, z305, z320 shown, full z-stack in Figure S14B. scale bar 100um. (C/D) annotations: a = amnion, ba = branchial arch, ce = corneal ectoderm, cme = cephalic mesoderm, da = dorsal aorta, de = diencephalon, drg = dorsal root ganglion, fb = forebrain, flb = forelimb bud, g = gut, h = heart, hb =. Hindbrain, hlb = hindlimb bud, hbw = hindbrain wall, hgd = hindgut diverticulum, itcd = internal carotid artery, l = lense, mb = midbrain, me = mesencephalon, mcv = mesencephalic vesicle, mes = mesonephros, mg = midgut, nc = notochord, nt = neural tube, o = oral cavity, oc = inner layer of optic cup, s = somites, sc = spinal cord, sv = sinus venosus, tb = tail bud, te =telencephalon. E. schematic of embryonic stages XXII-XXVI.

We then analysed the latest stage of pre-oviposition development (Figure 6D/S14A/B M225A). Here, we noticed the prominent midbrain growth, which we already had observed in the brightfield images (Figure 6A). This led to the first sections showing the mesencephalic vesicle formed by the neuroepithelium which is covered by the surface ectoderm (z000-056) with the cephalic mesenchyme in-between at deeper sections (z046-z056). Then, the optic vesicles appear (z072-150), which have developed into full cups with a lens in the middle overlaid by the surface ectoderm. The optic fissure remains open as a small slit (z111). Directly adjacent to the optic cups (z111-131), the olfactory placode is visible. The otic vesicles are clearly distinguishable (z137-174) and the heart exhibits all ventricles (z165-232) with blood cells present in the heart as well as in the main arteries and veins (z205-292). Continuing deeper into the embryo, the oral cavity becomes visible which then continues into the midgut (z195-z239). Forelimb– and hindlimbs can be distinguished (z200-308). Notably, the mesonephri appear very prominent between the heart and tail (z239-292). The somites adjacent to the neural tube/spinal chord are also clearly visible as square epithelialized structures on either side of the neural tube (z297-323).

**Stage XXII**: The 3^rd^ pharyngal arch is present, and the tail is elongated and curled. About 20 somites.

**Stage XXIII**: First signs of limb bud initiation. 22-25 somites.

**Stage XXIV**: 4^th^ pharyngal arch formation has begun, the optic fissure has narrowed, the tail elongated and the mesonephri are more prominent. 26-28 somites.

**Stage XXV:** The 4^th^ pharyngal arch is present, clear fore– and hind limb buds are present, and the optic fissure is very narrow. About 31 somites.

**Stage XXVI:** The 5^th^ pharyngal arch has begun to form, the forelimb and hindlimb buds continue to grow, and the tail continues to elongate. The optic fissure is almost closed, and eye pigmentation has begun. Up to 38 somites.

## Development of the Central Nervous System and Neural Crest Cell Migration

After establishing our staging system for pre-oviposition anole embryogenesis, we utilized it to explore the development of the central nervous system, neural crest cell migration, heart morphogenesis and muscle differentiation in more detail. For this, we co-stained embryos of consecutive stages with the neuronal marker TUJ1, neural crest cell marker HNK1 or myosin heavy chain marker MF20.

We initially focussed on the central nervous system starting with DAPI stained XVII stage embryos (Figure 7A). At this stage, TUJ1 expression was diffuse but could be detected in the head folds and the rostral frontonasal tip of the embryo (Figure 7B, stage XVII). TUJ1 staining then becomes more localised to the midbrain, hindbrain and trunk spinal cord (Figure 7B, stage XX, box). TUJ1 was also observed labelling axons perpendicular to the neural tube, along the dorsal side of the caudal somites (arrows). This pattern is consistent with early neuronal differentiating of emerging trunk neural crest cells. At stage XXI, TUJ1 becomes highly expressed in the brain, extending from the hindbrain to the optic vesicle and presumptive olfactory placode region, where it broadens into a delta shape (Figure 7B, stage XXI, box).. At stage (XXIIa/b), TUJ1 staining extends from the hindbrain towards the optic vesicle with the delta-like pattern in the presumptive olfactory region inferior to the optic vesicle persisting (Figure 7B, stage XXIIa box). TUJ1 expression becomes then more pronounced in the forebrain (Figure 7B, stage XXIIb, box ii) and along the neural tube. It also localises adjacent to the pharyngal arches, perhaps consistent with initiation of cranial nerve formation (Figure 7B, stage XXIIb, box i), while the metameric track like pattern of axons along the spinal cord begins to extend further towards the tail consistent with progressive formation of the neural crest cell derived peripheral nervous system. These patterns intensify with TUJ1 labelling extending into the pharyngal arches consistent with formation of cranial nerves (Figure 7B, stage XXIII, box i). The localized labelling of the olfactory vesicle becomes more distinct (Figure 7B, stage XXIII, box ii). At stage XXIV, these thin lines of TUJ1 labelled cells extending towards the pharyngal arches have become streams (Figure 7B, stage XXIV, box i) and the labelled area surrounding the olfactory vesicle is also much more pronounced (Figure 7B, stage XXIV, box ii). Furthermore, the patterns of presumptive migrating neural crest cells, along the neural tube as they begin to form the peripheral nervous system, has become more intense and the head contains a plethora of labelled future nerve fibres coursing throughout the brain.

**Figure 7:**
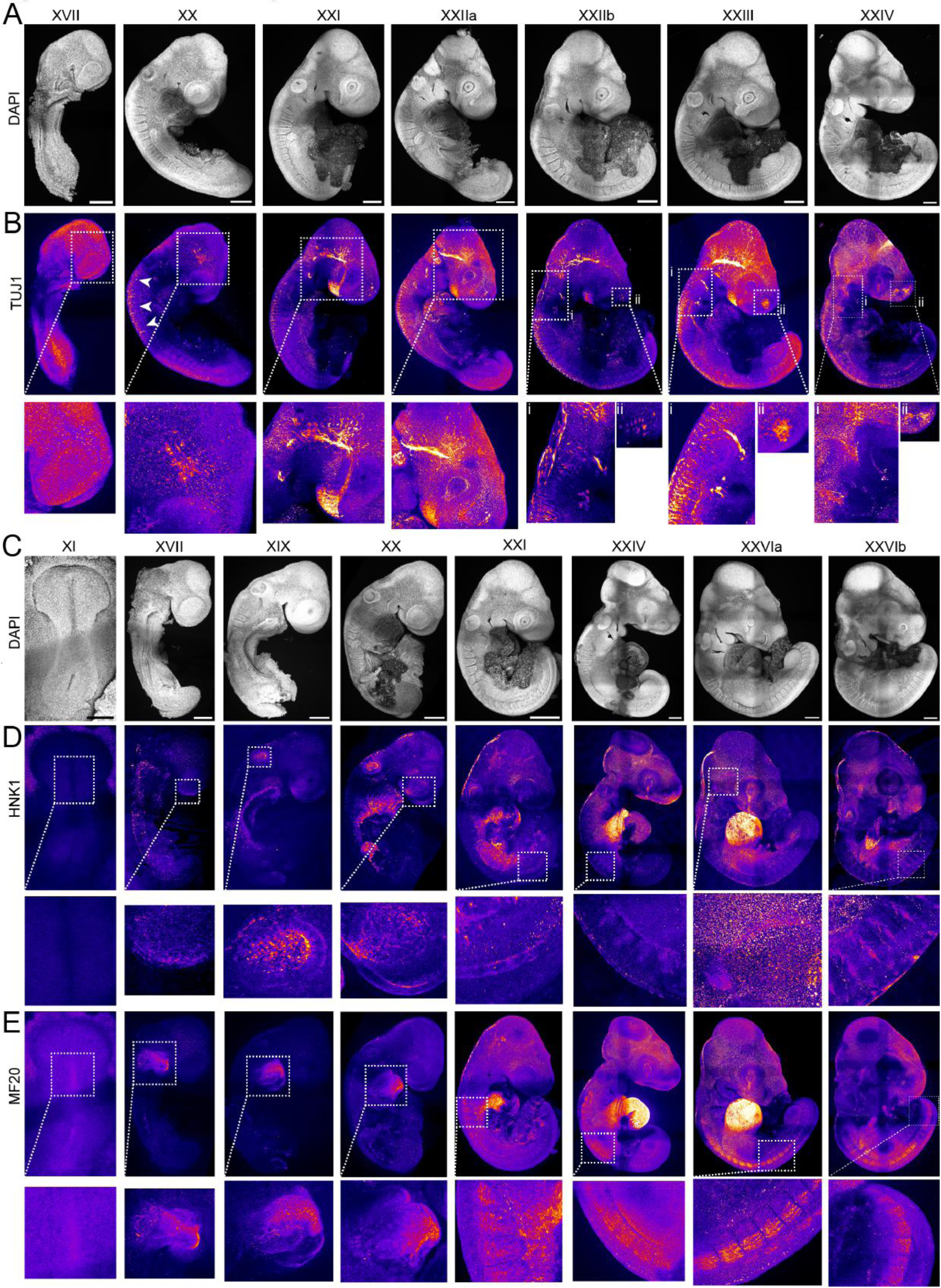
Neural Crest Cell Migration and Development of the Central Nervous System. **A-B.** Maximum Intensity Projections of Immunofluorescence stainings of embryos stages XVII-XXIV. Fire staining corresponds to signal intensity with yellow as high and purple as low signal. (D) DAPI, (E) central nervous system marker TUJ1. XVII=(M), XX=(M), XXI=(M), XXIIa=(M), XXIIb=(M), XXIII=(M), XXIV=(M). scale bars = 250um. (E) bottom row shows zoom in of white boxed regions of top row. Fire staining corresponds to signal intensity with yellow as high and purple as low signal. **D-E.** Maximum Intensity Projections of Immunofluorescence staining of embryos stages XI-XXVI (A) DAPI, (B) myosin heavy chain marker MF20, (C) neural crest cell marker HNK1. Embryos stage XI=(M149B), XVII=(M219B), XIX=(M206B), XX=(M230A), XXI=(M207A), XXIV=(M222A), XXVIa=(M216A), XXVIb=(M206A). scale bars=250um. (B/C) bottom rows zoom-ins of white box in top rows.

We next investigated the migration of neural crest cells by analysing the expression of HNK1 (Figure 7C,D). At stage XI, during early neurulation, the entire embryo is negative for HNK1, suggesting that there are no migrating neural crest cells present at this stage (Figure 7D, stage XI, white box). At stage XVII, HNK1 labels the rostral tip of the embryo, the forebrain (Figure 7D, stage XVII, white boxes), and can be detected in the optic vesicle, hindbrain, and otic vesicle, (Figure 7D, stage XIX). HNK1 then continues to be highly expressed in the forebrain, optic and otic vesicles, and can be found at lower levels in cells spread throughout the cranial mesenchyme in the developing head (Figure 7D, stage XX). At stage XXI, HNK1 expression continues in the hindbrain, otic vesicles and the optic vesicles, but not in the lens or in streams covering the head (Figure 7D, stage XXI). Subsequently at stage XXIV, HNK1 labels neural crest cells migrating through the anterior half or each somite, which is a highly conserved pattern in vertebrates. HNK1 also labels streams of presumptive neural crest cells in the mesenchyme covering the brain but excluding the mesencephalic vesicle. HNK1 is also expressed inside the optic vesicle and prominently labels the optic fissure, as well as hindbrain and presumptive neural crest cells (Figure 7D, stage XXIV). At the final stage of preoviposition development (stage XXVI), HNK1 continues to label migrating neural crest cells throughout the mesenchyme surrounding the brain, and within the anterior half of each somite.

To gain insight into heart and muscle development, we analysed MF20 expression (Figure 7E). MF20 can be found distinctively in the upper half of the embryonic disk, close to the head folds, demarcating the position of the future heart already at stage XI (Figure 7E, stage XI, white box). At stage XVII, MF20 is enriched caudal to the head, revealing the initiation of heart morphogenesis. The heart then enlarges and becomes a clear sac with inner looping (Figure 7E, stage XIX, box and arrows). At stage XX, MF20 demarcates the heart (Figure 7E stage XX). Thereafter, MF20 continues to he expressed in the heart but shortly afterwards is also detected in the presumptive myotomes of the first 8 somites, as well as in presumptive myoblast in the head and other cells in the lens (Figure 7E, stage XXI). Subsequently at stage XXIV, MF20 is expressed in more posterior somites. and continues to label presumptive cranial myoblast as well as the inner surface of the optic vesicle (Figure 7E, stage XXIV). At stage XXVI, MF20 exhibits high persistent expression in the heart, and demarcates presumptive striated myoblasts in the maturing dorsal dermomyotome of the somites. (Figure 7E, stage XXVIa/b). MF20 continues to label cranial myoblasts as well as the presumptive myogenic cores of the pharyngal arches.^42–44^

## Discussion

This work provides the first study of *A. sagrei* pre-oviposition development describing 26 distinct stages of embryogenesis from fertilisation up to limb bud formation and eye pigmentation that are characterised by brightfield imaging, transmission electron microscopy, tissue sectioning, histology and immunostaining. Our analysis of the ovarian follicles in *A. sagrei* is the first for this species. We find that, similar to other reptiles, anole oocytes contain a germinal vesicle of about 150μm, which is surrounded by lipid and yolk vesicles as well as mitochondria and other subcellular structures.^32,33,45^ While the germinal vesicle has been generally described in other reptilian species, very few recent studies have focused on the architecture this structure and, to our knowledge, this structure has never been described in the brown anole. At this stage, the chromosomes are supposed to be diplotene but further studies are required to confirm this for anoles.^46^ We were unable to detect any spindle apparatus which raises the question of how meiosis occurs and how the mechanics of the first cleavage divisions are mediated. Similarly, we do not yet understand how ovulation or fertilisation occur in this species. We know ovulation to be hormonally controlled in the genus and that the cellular architecture of the follicle wall changes following hormonal changes in preovulatory female anoles.^47^ However, the process of rupturing the single cell layer surrounding each follicle, the process of fertilisation, and follicular dynamics after ovulation have not yet been described in this species. Further studies are therefore required to shed light on these biological processes.

The cleavage division stages appear conserved across avian and non-avian reptiles with large furrows, and blastomeres, which then divide into smaller and smaller blastomeres as the cleavage divisions continue (Figure 2 B/C).^2,5^ Interestingly, we could observe furrows in embryos that already consisted of a high number of cells with several cells forming the furrows (Figure 2D), which has not been reported in chicken or *Zootoca*.^2,5^ This could also be due to the fact that these previous studies relied solely on brightfield images, permitting less detailed analysis. Further analyses will therefore be required to understand whether this feature is specific to anoles or whether it is conserved across other clades.

We found that peri-gastrulation development in anoles significantly diverges from chicken and *Zootoca*. While the chicken embryo is flat and monolayered with a hypoblast that migrates ventrally, the anole is multilayered and adopts a dome-like shape. In contrast, *Zootoca* exhibits a tunnel-like morphology. Notably, while the anole embryo directly overlays the yolk, the chicken embryo is stretched over the subgerminal cavity, a lumen.^3,14^ We also observed that the slit-like blastopore in anoles is oriented in parallel to the anterior-posterior axis while it is perpendicular in *Zootoca* which could be indicative of different tissue mechanics driving the ingression process.^5^ Collectively, these data suggest that contrary to current opinion, chicken morphogenesis is not representative of all avian and non-avian reptile embryogenesis.^48^ Further studies of representatives of different clades are therefore required to comprehensively analyse similarities and differences and to understand whether these results are just exceptions. However, we believe reptile embryogenesis in general to be more diverse than previously thought or appreciated.

The anole embryo remains highly curved during the initiation of neurulation and only flattens once the neural tube closes, while the chicken embryo remains flat until the head rotates.^3^ We could observe only one closure point during anole neurulation, which is similar to chicken embryogenesis^49^ but different from mammalian neurulation, which exhibits several neural tube closure initiation points.^50^ Following neurulation, the anole appears much more similar to chick and *Zootoca* which is in accordance with the hourglass model of development. This model describes the divergence of all species within one phylum during early embryogenesis, their convergence during organogenesis, and their divergence again upon final body plan acquisition.^51^ However, it will require more detailed cross-species analysis to understand the mechanistic basis of these conserved and divergent features.

Our staging series enables comparative analyses of pre-oviposition *A. sagrei* embryos with other squamates as well as cross-clade analyses with avian reptiles or mammals. Our analysis of early neuromuscular development through localisation of the neuronal marker TUJ1, neural crest cell marker HNK1, and muscle-marker MF20 can be used for cross species comparison with recently published studies in the veiled chameleon^40^ and the common wall lizard^52^. The divergence of peri-gastrulation morphogenesis between *Zootoca*, chicken and anoles warrants further in-depth analyses of the precise differences, and may imply that morphological differences affect tissue mechanics and be a sign of possible differences in signalling pathway dynamics. For example, it would be highly interesting to investigate the process of neurulation in more detail, where both chicken and *Zootoca* have a straight midline while the anole exhibits a highly convex midline, which may result in different forces being required for elevation of the neural folds. Furthermore, it will be interesting to compare the timing of organogenesis to understand whether different rates of development exist for different organs and what implications such heterochronies have for subsequent development, and whether they may reflect different requirements for hatchlings/neonates.

The elucidation of developmental stages for the brown anole lays the foundations of a higher resolution picture of comparative developmental biology that compares and contrasts the features of cellular processes and mechanisms in evolutionarily related and more distant organisms. Our understanding of developmental biology, cellular processes and mechanisms is rooted in a limited set of model organisms that are considered representative for their respective group or even class. As such, the mouse is commonly used to broadly model mammalian embryogenesis. Similarly, the chicken has been used to represent all avian and non-avian reptiles^14^ despite the fact that proper detailed comparative analyses between chick and other non-avian reptile species have not yet been conducted to substantiate this philosophy. Therefore, we do not yet understand how much developmental variation exists among closely related species or among the major vertebrate radiations. This is due in part to a to dearth of non-traditional model organisms and the lack of databases compared to human or mouse development.^53,54^ Our analysis here suggests that while early cleavage divisions appear conserved, the chicken does not accurately represent non-avian reptile species peri-gastrulation and early neurulation morphogenesis. Further in-depth studies using higher resolution microscopy are therefore required to better understand the mechanisms underpinning the variations at these later developmental time points. Our morphological staging analyses of brown anole development should now be complemented by a molecular characterisation of embryogenesis, including investigation of transcriptional programs and their functional significance. Through these approaches we can continue to derive important evolutionary and developmental insights from non-traditional model organisms.

## Acknowledgements

The authors would like to thank all members of the Trainor lab and Dr Suzannah Williams for critical feedback. The authors would like to thank Sean McKinney for his support processing the confocal images. Research in the Trainor lab is supported by the Stowers Institute for Medical Research, and N.A.S is supported by a K99 Pathway to Independence award from the National Institute for Child Health and Development (HD114881). The Sanger lab is funded by the USA National Science Foundation (#1942250). AW is supported by a postdoctoral research fellowship of All Souls College, University of Oxford, and received a travel grant of the Cambridge Philosophical Society. AW and FH were supported by the Newton Trust, the European Research Council (695669) and the Engineering and Physical Sciences Research Council (EP/Y032756/1). Upon publication, original data underlying this manuscript can be accessed from the Stowers Original Data Repository at http://www.stowers.org/research/publications.

## Author Contributions

AW conceived, planned and carried out the project. NAS supported project planning and protocol optimisation. BKK annotated H&E cross sections. HW carried out paraffin sectioning and Histology staining. MMC performed SEM and STEM. AW and MM dissected *A. sagrei* embryos. TJS and KS collected *A. sagrei* females. FH provided critical feedback and supervised AW. PT supervised the project. AW wrote the manuscript with the help of TJS, NAS, BKK, FH, and PT.

## Declaration of interests

The authors declare no competing interests.

## Methods

### Collection of females

All protocols described were approved by the Loyola University Chicago IACUC committee (#3662). Gravid females of the brown anole (*A. sagrei*) were collected in Miami, FL, from May 2^nd^-10^th^ 2023 according to State regulations. The females were housed in cloth bags for 24-96h until dissection.

Prior to dissection, the females were placed in a cooling box filled with one icepack for 5-10min. Snout-to-vent length was measured from head to vent prior to euthanasia.

## Dissection

Following euthanasia, the female was placed on its back and its abdomen opened from the cloaca through to its sternum. The female reproductive tract was dissected away from the body into phosphate-buffered saline (PBS). The follicles were then dissected from the ovaries, otherwise, the reproductive tract was fixed for further analysis or discarded.

## Follicle Isolation

Yolk-filled follicles were isolated from clear follicles in the ovaries and the ovaries were fixed in 4% PFA. In yellow yolk-filled follicles, the germinal vesicle could be distinguished within the germinal disc, a lighter-coloured, round structure on the surface of the oocyte. In the middle of the germinal disc, a round, dark shadow could be defined which is the germinal vesicle. To isolate the germinal vesicle, the follicles were oriented such that the germinal disc was facing up. Then, the germinal disc was isolated with an underlying layer of yolk. The germinal vesicle could be fully isolated from the germinal disc but was then very sticky and often lost following fixation. It proved more feasible to fix the germinal disc with the germinal vesicle inside and then dissect the germinal vesicle from the fixed structure if need be. Here, the germinal discs were fixed in 4% PFA and the germinal vesicles further dissected.

## Embryo Isolation

All embryo dissections were carried out in PBS. The eggshell was held in place by forceps and a small incision was made using Vannas Spring Scissors (Fine Science Tools 15000-03). Then, this incision was used to grip the eggshell with forceps and the egg was cut open along the long axis starting at the incision point. It is important to angle the scissors in parallel with the eggshell and avoid cutting deep into the egg as this can damage the embryo. Once the eggshell was cut in 2 halves, the eggshell was carefully removed from the yolk. The embryo is located on the long side of the egg, so the egg could carefully be turned using the two ends as anchor points. Early-stage embryos could be localised through the position of the germinal disc that appears almost white on top of the yellow yolk and is easily recognised.

**Cleavage stage** embryos are not cohesive structures, so they must be dissected with a layer of yolk beneath them to remain intact. For this, the germinal disc was carefully cut out of the embryo, cutting through the yolk to retain the entire disc intact. A needle was then used to cut through the yolk beneath the germinal disk leaving about 1-2mm of yolk underlying the germinal disk for support. It is important to keep the disk during this process as flat as possible, otherwise the embryo will rip along the cleavage furrows.

**Peri-gastrulation stage** embryos are cohesive and can also be localised by determining the position of the germinal disk. The embryo is in the middle of the germinal disk; thus, it could be dissected by cutting along the border of the germinal disk. Once that was complete, the embryo could detach on its own from the yolk. If that was not the case, forceps were used to lift the germinal disk carefully from the yolk. Then, a needle was used to cut through attachments between embryo and yolk while the embryo is continuously peeled off. The embryo did not constitute the entire germinal disk but could be found as a dome-like structure in the middle of the disk. Once removed from the yolk, the surplus tissue could be removed either by scissors or two needles.

During **early neurulation stages**, the blastoderm has grown around the entire egg and thus its edge cannot be used anymore to locate the embryo. Instead, the embryo is found at the opposite side of the blastoderm closure. The egg was turned from the side until the embryo, which is visible with a distinct shape (neural tube and beginning head folds) on top of the yolk but has the same colour as the rest of the egg, faced upwards. The embryo was cut out of the blastoderm using scissors leaving 1-2mm of blastoderm around the embryo. In case the embryo did not separate from the yolk on its own, and forceps were used to detach the embryo from the yolk. Then, the embryo was cleaned using needles. First, the surrounding blastoderm was cut off and then the amnion cleared away.

For **late neurulation and organogenesis stages**, the embryo can readily be distinguished once the eggshell was removed as it is located in the middle of blood vessels. First, the embryo inside the chorion was separated from the surrounding yolk using scissors and forceps. Then, using forceps and a needle, the chorion was opened and – in case the embryo did not move out of the chorion on its own – cut further open along the dorsal side of the embryo. Once the chorion was removed, the embryo was still enveloped by the amnion, which lies tightly over the entire developing body. The amnion was pierced by a needle between the forebrain and the first pharyngal arch, then it was held in position by forceps, while a needle was used to open the amnion around the head and then along the back up to the tail bud.

## Fixation and embryo storage

Following dissection, each embryo was pipetted into a 1.5ml Eppendorf tube filled with 1ml 4% PFA and fixed at 4°C overnight or moved into a 4 ml glass vial and fixed in glutaraldehyde-PFA solution pH 7.4.

Embryos fixed in PFA were then washed 3x with PBS and then dehydrated. Embryos allocated for histology were dehydrated stepwise into 70% EtOH (5min each in 25%, 50%, 70%) and stored at 4°C. Embryos allocated for immunofluorescence analysis were dehydrated into 100% MeOH (5 min each in 25% MeOH, 50% MeOH, 75% MeOH, 95% MeOH, 100% MeOH wash, 100% MeOH) and stored at –20°C until required for analysis. Embryos allocated for SEM were kept in glutaraldehyde-PFA solution.

## Brightfield imaging

Embryos were placed in a petri-dish in either 70% EtOH, 100% MeOH or Glutaraldehyde and imaged on a Leica DFC550 microscope through manual stacks. The stacks were rendered using Helicon (Focus 5.3 software). Each embryo was imaged from its dorsal – and ventral (pre-gastrulation to Neurulation) or left – and right side (neurulation-organogenesis).

## Paraffin cross sections and Histology

Fixed embryos were paraffin processed on a Milestone Pathos Delta microwave tissue processor without pressure to preserve embryo morphology. Embryos were embedded in paraffin wax and sectioned at 4µm onto charged slides coated with Poly-L-Lysine. Slides were baked at 60C for at least an hour before standard H&E staining to ensure attachment of delicate structures

## Electron microscopy

Samples were fixed in 2.5% glutaraldehyde and 2% paraformaldehyde. For SEM imaging, a TOTO processing strategy was used (reference 1) with 1% aqueous tannic acid, 1% osmium tetroxide and 2% thiocarbohydrazide (TCH), followed by dehydration in a graded series of ethanol and critical point drying in a Tousimis Samdri 795. Samples were mounted on stubs and coated with 4nm gold palladium in a Leica ACE600 coater before imaging in a Zeiss Merlin SEM at 8kV with SE2 or InLens detectors.

Samples prepared for sectioning and STEM imaging were fixed as for SEM, followed by secondary fixation in 1% buffered osmium tetroxide and *en bloc* staining with 0.5% uranyl acetate. Samples were then dehydrated in a graded series of ethanol with propylene oxide as a transitional solvent and infiltrated with a graded series of Hard Plus resin (Electron Microscopy Sciences) and propylene oxide. After embedding and polymerization at 60°C for 48 hours, blocks were trimmed of excess resin and imaged with a Bruker SkyScan 1272 microCT at 4 watts, with pixel sizes ranging from 0.68um to 3um at either 50kV with no filter or 60kV with Al 0.25mm filter, using 0.1-0.2 degree steps and 180 degree imaging. MicroCT volumes were used to target the center of nuclei for sectioning at 80nm on slot grids using a Leica UC7 ultramicrotome and Diatome diamond knife. Sections were post stained for 6 minutes each with 4% uranyl acetate in 70% methanol and Sato’s triple lead stain, and imaged in a Zeiss Merlin SEM with aSTEM detector at 26kV and 700pA.

## Immunostaining

Mid-neurulation to organogenesis stage embryos were bleached for 2h using Dent’s bleach (Hydrogen Peroxide, DMSO, MeOH (1:1:4)) followed by 2 washes in 100% MeOH. All embryos, follicles and germinal vesicles were then re-hydrated into PBS (75%, 50%, 25%, 0% MeOH 5min each), washed 2x in PBS and then permeabilised for 20min (early stage embryos) to 2h (organogenesis stages) in permeabilization buffer (0.3% Triton X-100, 0.1 M Glycin in PBS). The samples were blocked for 1h at RT in blocking solution (PBS, 0.1% Triton X-100, 1% v/v donkey serum) and incubated in primary antibodies overnight at 4°C on a rocker. The samples were then washed 3x for 10min in PBST (PBS, 0.1% Triton X-100) and incubated in secondary antibodies and DAPI for 4h at RT in the dark or overnight at 4°C. The samples were then washed 3x 10min in PBST and equilibrated through a glycerol series (25%, 50% glycerol in PBST for 15min each). The samples were then mounted in VECTASHIELD mounting medium (Vector Laboratories H-1200-10) between two cover slips using vacuum grease to enable imaging from both sides on a Leica SP8 confocal microscope with a 20x air objective using tile scanning with 15% stitching. The nucleoli of germinal vesicles were imaged on a Nikon Eclipse Ti2 microscope equipped with a Yokagawa CSU W1 10,000 rpm Spinning Disk Confocal with 50um pinholes using a 100x oil objective.

Primary antibodies: HNK1 (mouse-IgM, Developmental Studies Hybridoma Bank, 1:50), MF20 (mouse-IgG, Developmental Studies Hybridoma Bank, 1:50), TUJ1 (mouse-IgG, Developmental Studies Hybridoma Bank, 1:50).

Secondary antibodies: DAPI, donkey-anti-mouse-IgM-546, donkey-anti-mouse-IgG-488.

## Image processing

### SP8

Imaging was performed using bidirectional scanning. Due to a misalignment of the two scanners, the images were corrected computationally by shifting every odd scan-line by 4 pixels to align with the even lines. This correction was applied prior to stitching in cases with multiple tiles. The simple correction and stitching were applied in python and ImageJ respectively.

The following embryos were not corrected as artefacts were introduced through the processing: Figure 3D M164B ventral, Figure 4 D M243B.

### DAPI staining

Pseudo-SEM images were obtained through a maximum intensity projection of confocal images of DAPI stained embryos as previously described.^38,39^ Fire intensity staining (yellow-white: high signal intensity, purple-black low signal intensity) which provides information on the thickness and density of the respective tissues in maximum intensity projections.

## Supplementary Figures

**Figure S1:**
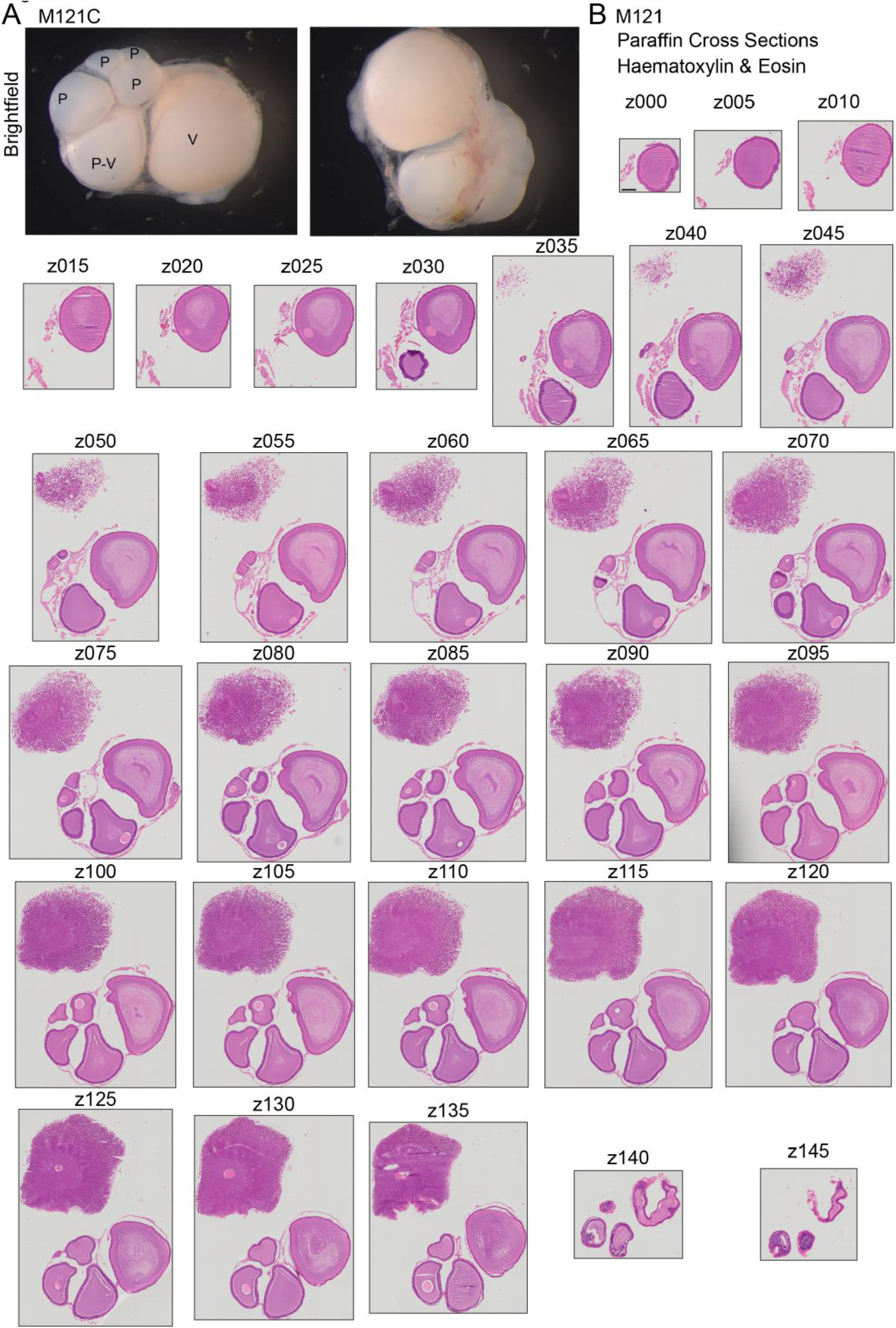
Morphology of maturing follicles. **A**. Brightfield images of follicles M121C. no scale bar. **B.** Hematoxylin and Eosin staining of paraffin cross sections of follicles M121C. Cross section number annotated in figure (z000-z145). scale bar: 250um. all images have the same scale.

**Figure S2:**
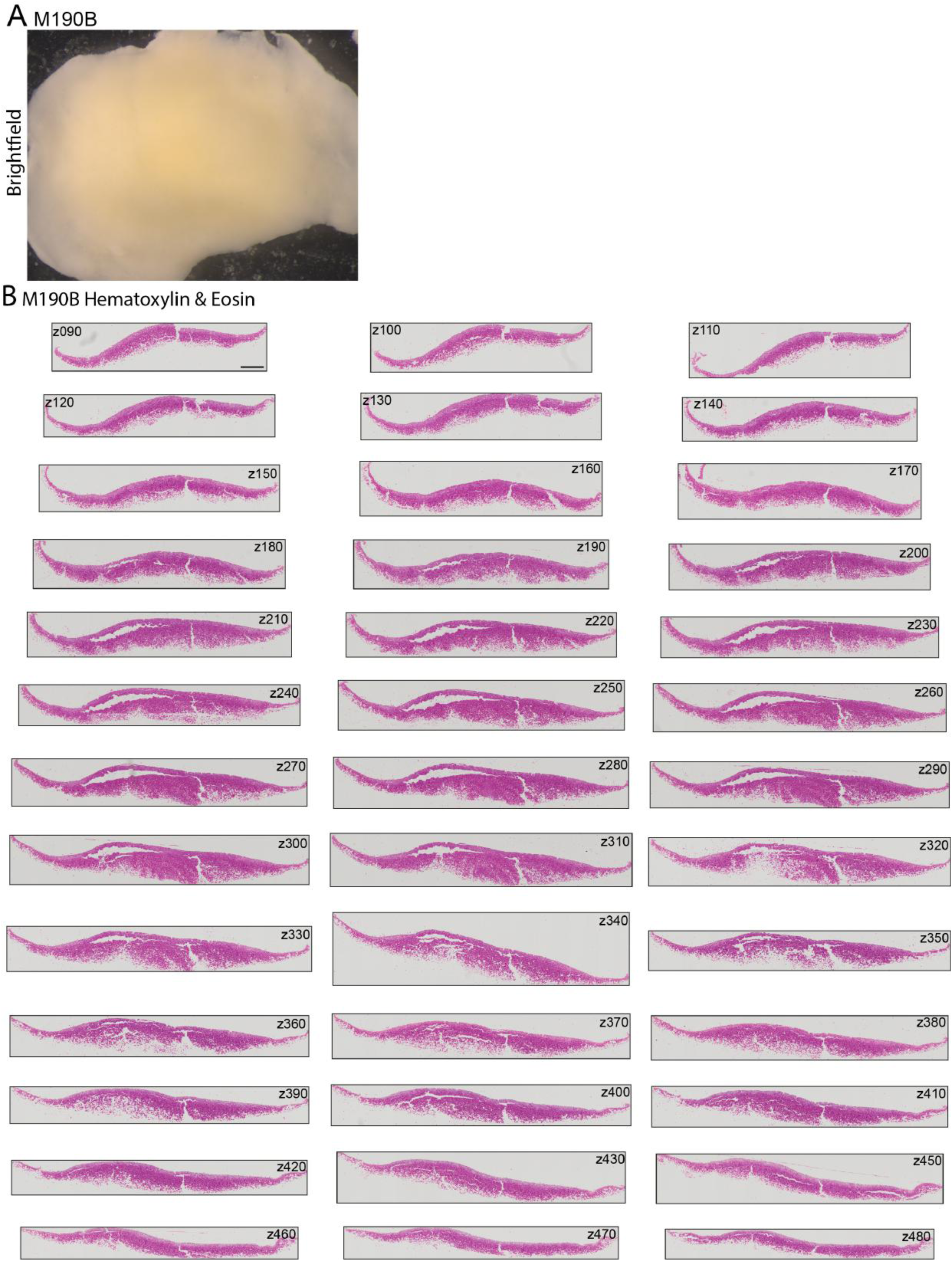
Cross sections of cleavage stage M190B. **A.** Brightfield image of M190B dorsal view. No scale bar. **B.** Hematoxylin and Eosin staining of paraffin cross sections of M190B. Cross section number annotated in figure (z090-z480). Scale bar: 400um. All images have the same scale.

**Figure S3:**
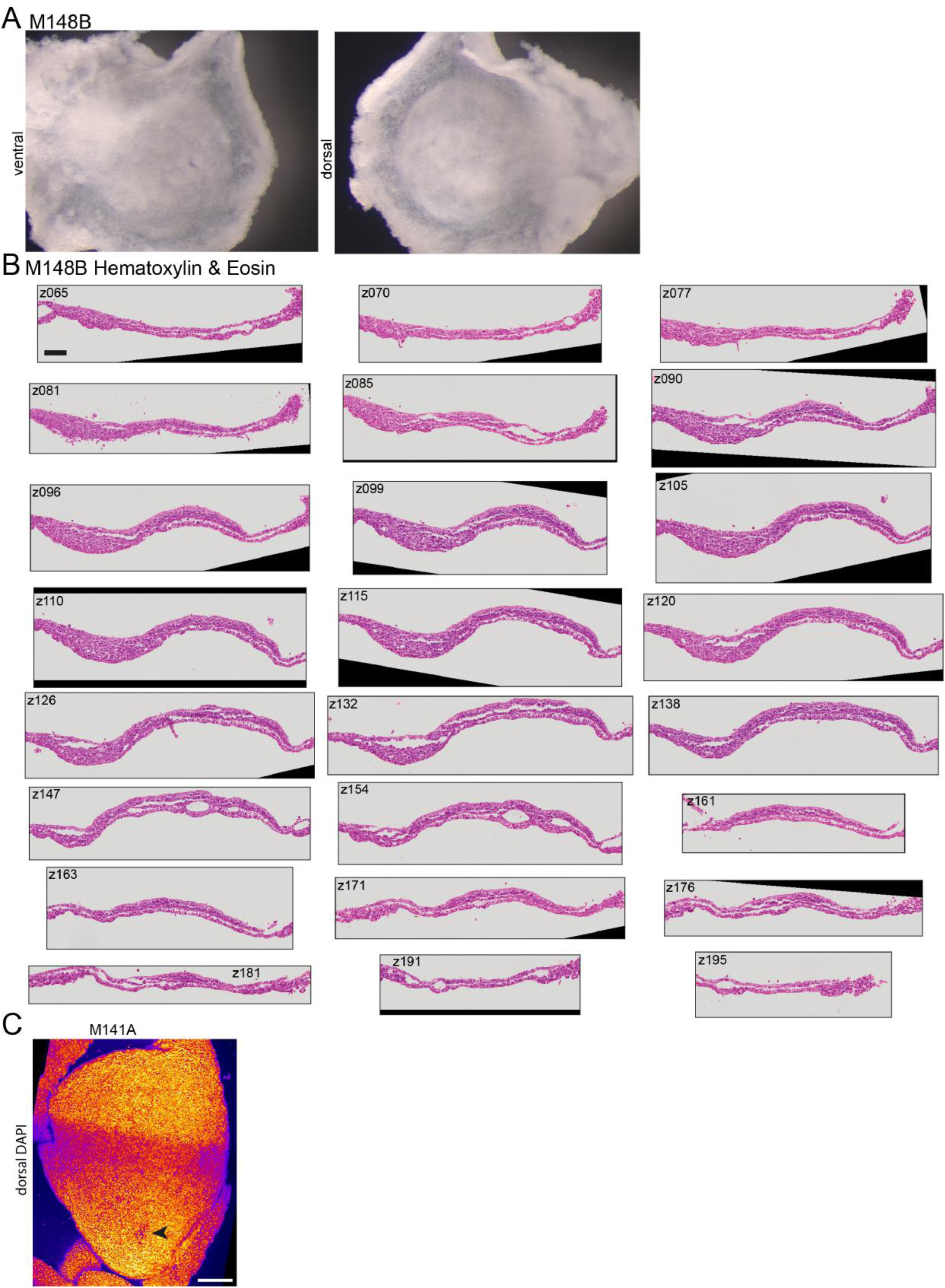
Cross sections of gastrulation stage embryo M148B. **A.** Brightfield images of M148B ventral and dorsal views. No scale bar. **B.** Hematoxylin and Eosin staining of paraffin cross sections of M148B. Cross section number annotated in figure (z065-z195). Scale bar: 100um. All images have the same scale. **C.** maximum intensity projection DAPI staining of M141A. blastopore is indicated with arrow. Scale bar 250um.

**Figure S4:**
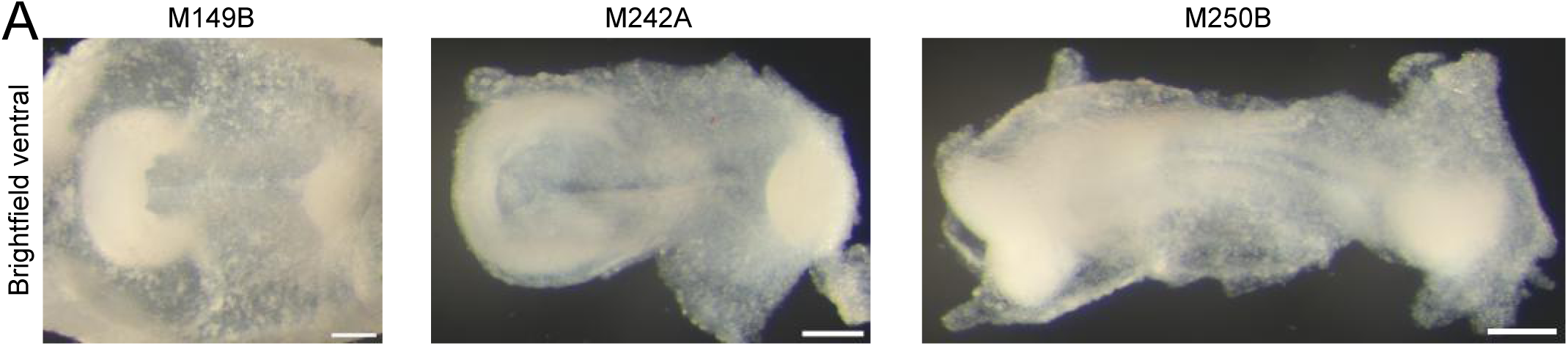
Brightfield images of initiation of neurulation ventral views. **A.** Brightfield images of ventral view of M149B, M242A, M250B. scale bars 200um.

**Figure S5:**
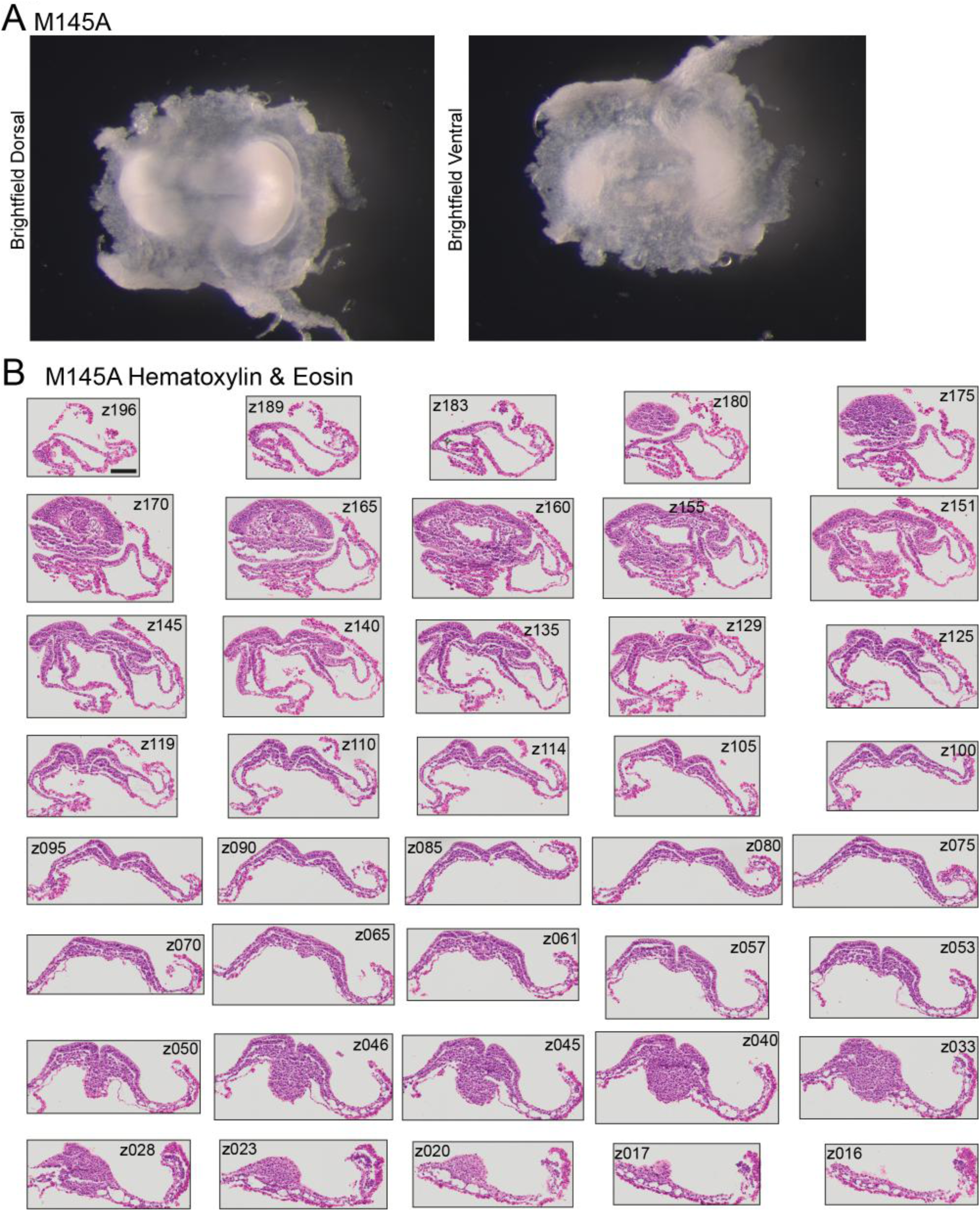
Cross section of initiation of neurulation M145A. **A.** Brightfield images of dorsal and ventral view of M145A. no scale bar. **B.** Hematoxylin and Eosin staining of paraffin cross sections of M145A. Cross section number annotated in figure (z196-z016 anterior to posterior). Scale bars 100um. All images have the same scale.

**Figure S6:**
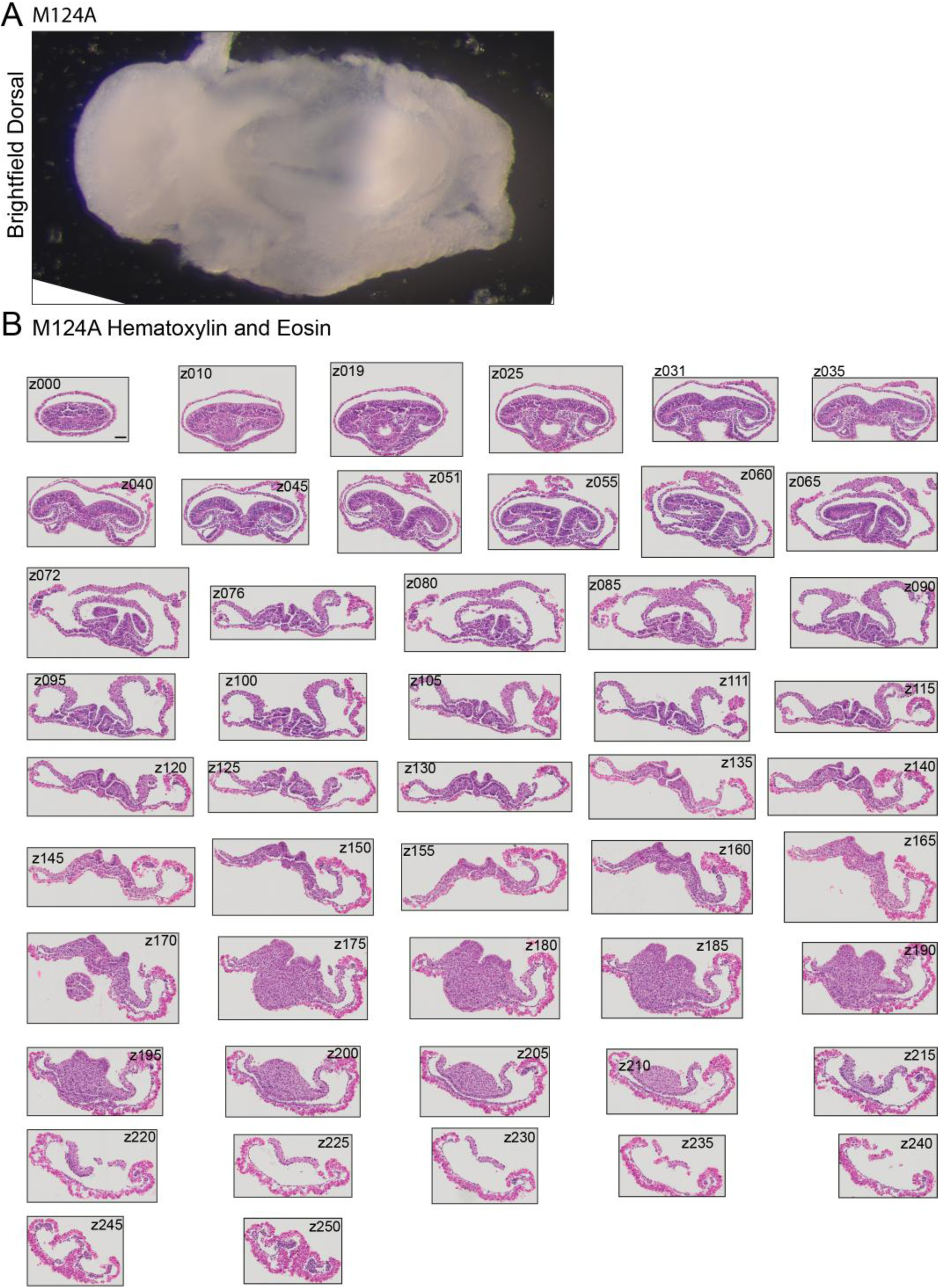
Cross sections of early neurulation embryo M124A. **A.** Brightfield image of dorsal view of M124A. no scale bar. **B.** Hematoxylin and Eosin staining of paraffin cross sections of M124A. Cross section number annotated in figure (z000-z250). Scale bars 100um. All images have the same scale bar.

**Figure S7:**
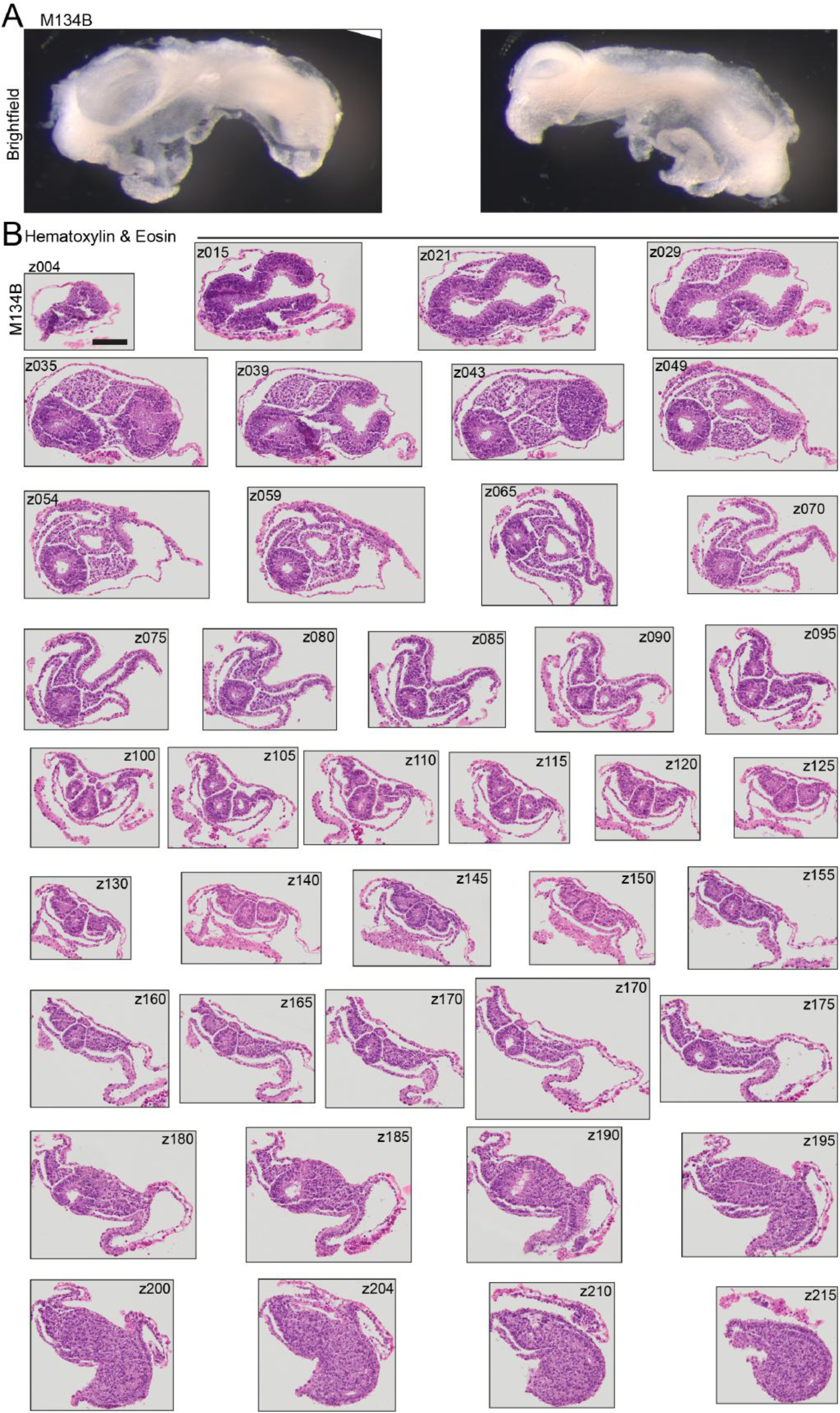
Cross section of neurulation embryo M134B. **A.** Brightfield image of right and left side of M134B. no scale bar. **B.** Hematoxylin and Eosin staining of paraffin cross sections of M134B. Cross section number annotated in figure (z004-z215). Scale bars: 100um. All images have the same scale bar.

**Figure S8:**
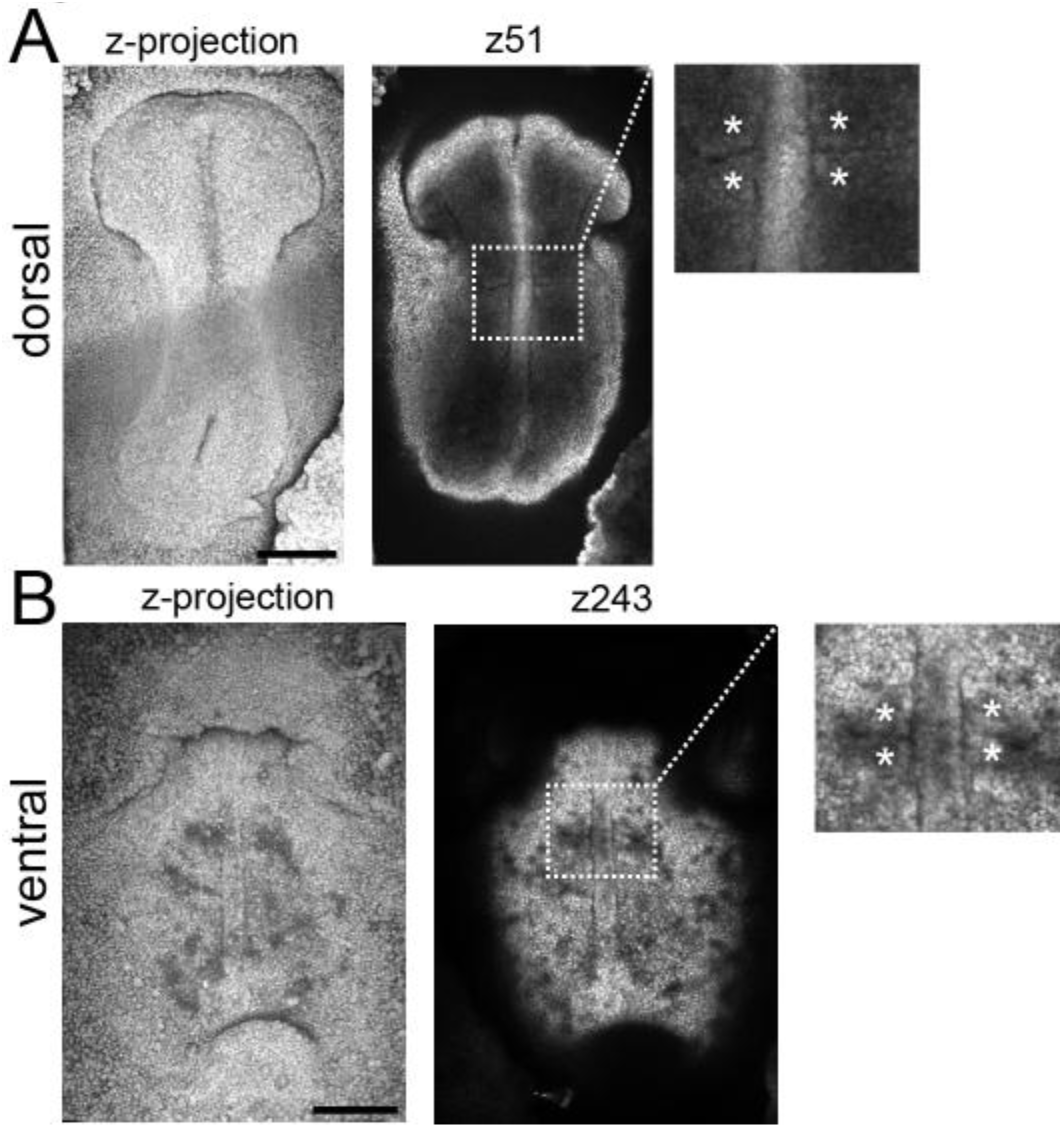
Somite Formation. DAPI stainings of M149B. **A.** M149B dorsal view. Left maximum intensity projection. Right: z51, somites boxed, asterisks in zoom-in demarcate somites. Scale bar: 250um. **B.** M149B ventral view. Left maximum intensity projection. Right z243, somites boxed. asterisks in zoom-in demarcate somites. Scale bar: 250um

**Figure S9:**
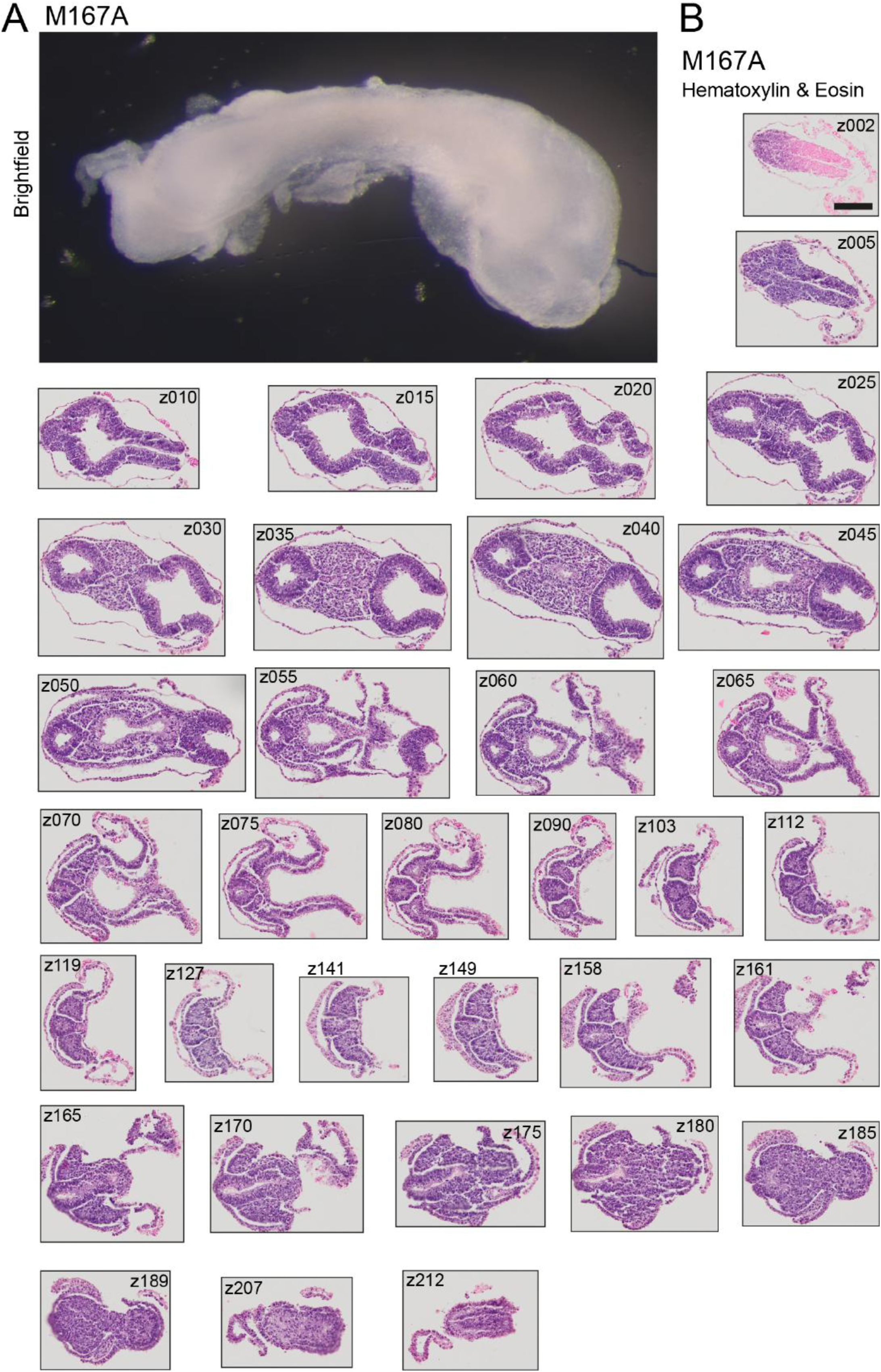
Cross sections of neurulation embryo M167A. **A.** Brightfield image of dorsal-right view of M167A. no scale bar. **B.** Hematoxylin and Eosin staining of paraffin cross sections of M167A. Cross section number is annotated in figure (z002-z212). Scale bar: 100um. All imaged have the same scale bar.

**Figure S10:**
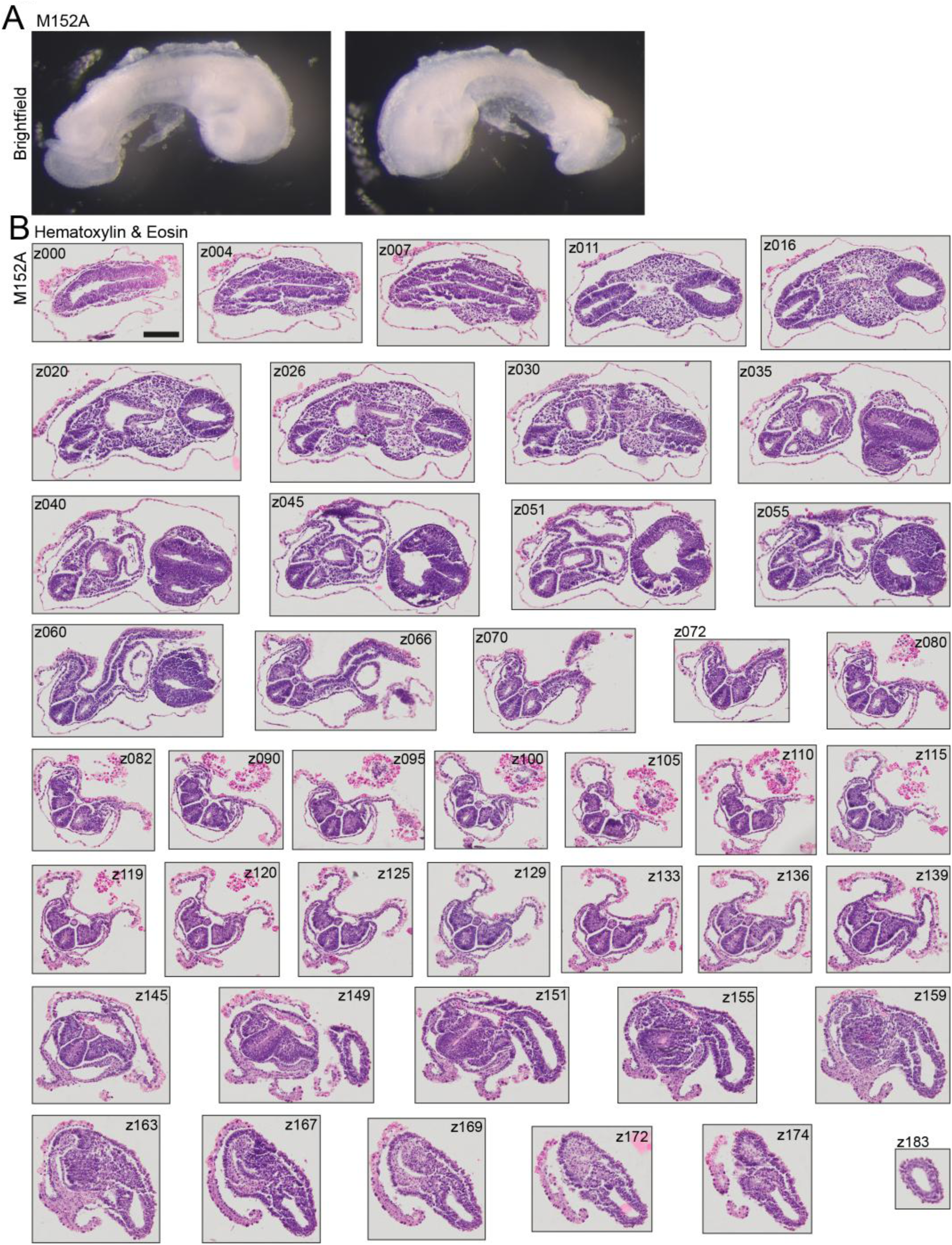
Cross sections of late neurulation embryo M152A. **A.** Brightfield image of right and left side of M152A. no scale bar. **B.** Hematoxylin and Eosin staining of paraffin cross sections of M152A. Cross section number is annotated in figure (z000-z183). Scale bar: 100um. All imaged have the same scale bar.

**Figure S11:**
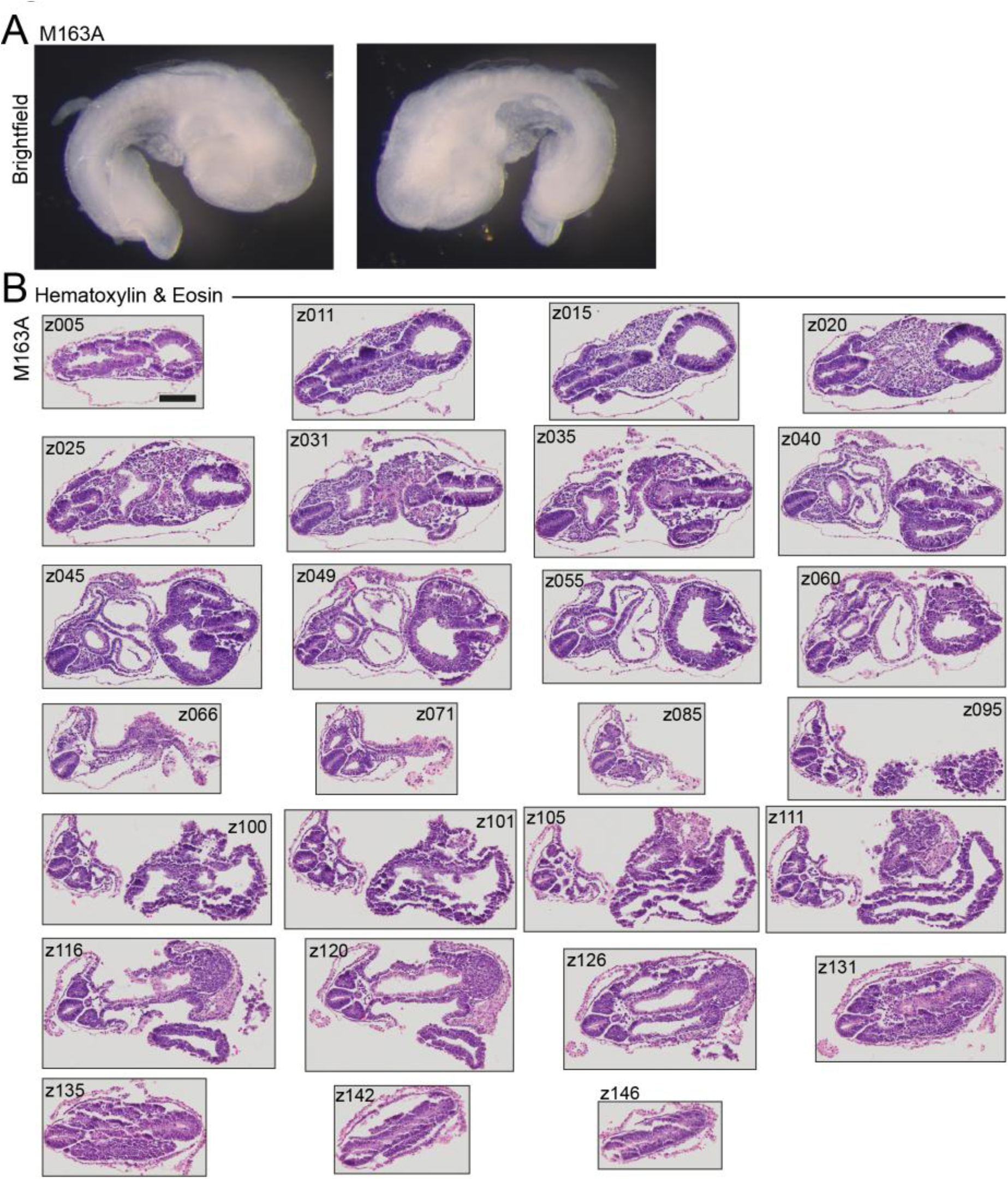
Cross sections of late neurulation embryo M163A. **A.** Brightfield images of right and left side of M163A. no scale bar. **B.** Hematoxylin and Eosin staining of paraffin cross sections of M163A. Cross section number is annotated in figure (z005-z146). Scale bar: 100um. All imaged have the same scale bar.

**Figure S12:**
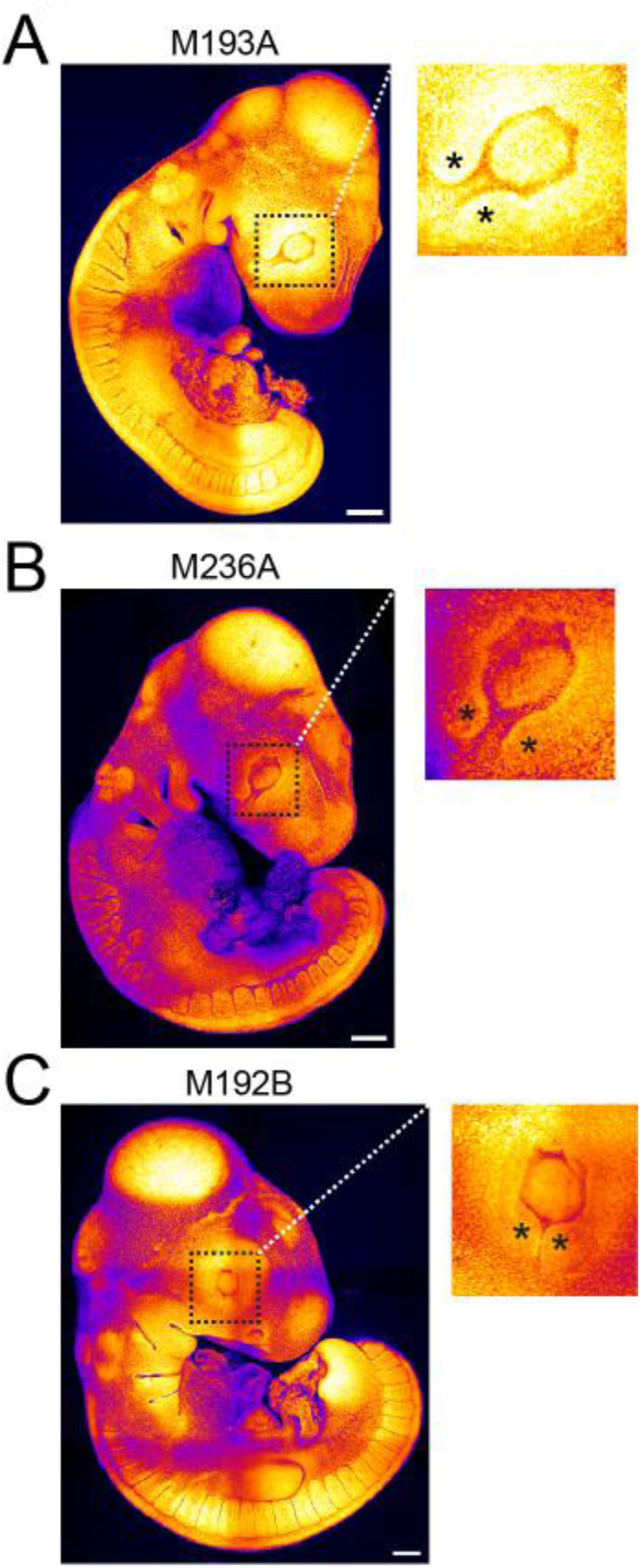
Closure of the optic fissure. **A-C.** Maximum intensity projections of DAPI staining of embryos (A) M193A, (B) M236A, (C) M192B. boxes around optical fissure. Asterisks in zoom ins (right column) demarcate the edges of the optical fissure. Scale bars 200um.

**Figure S13:**
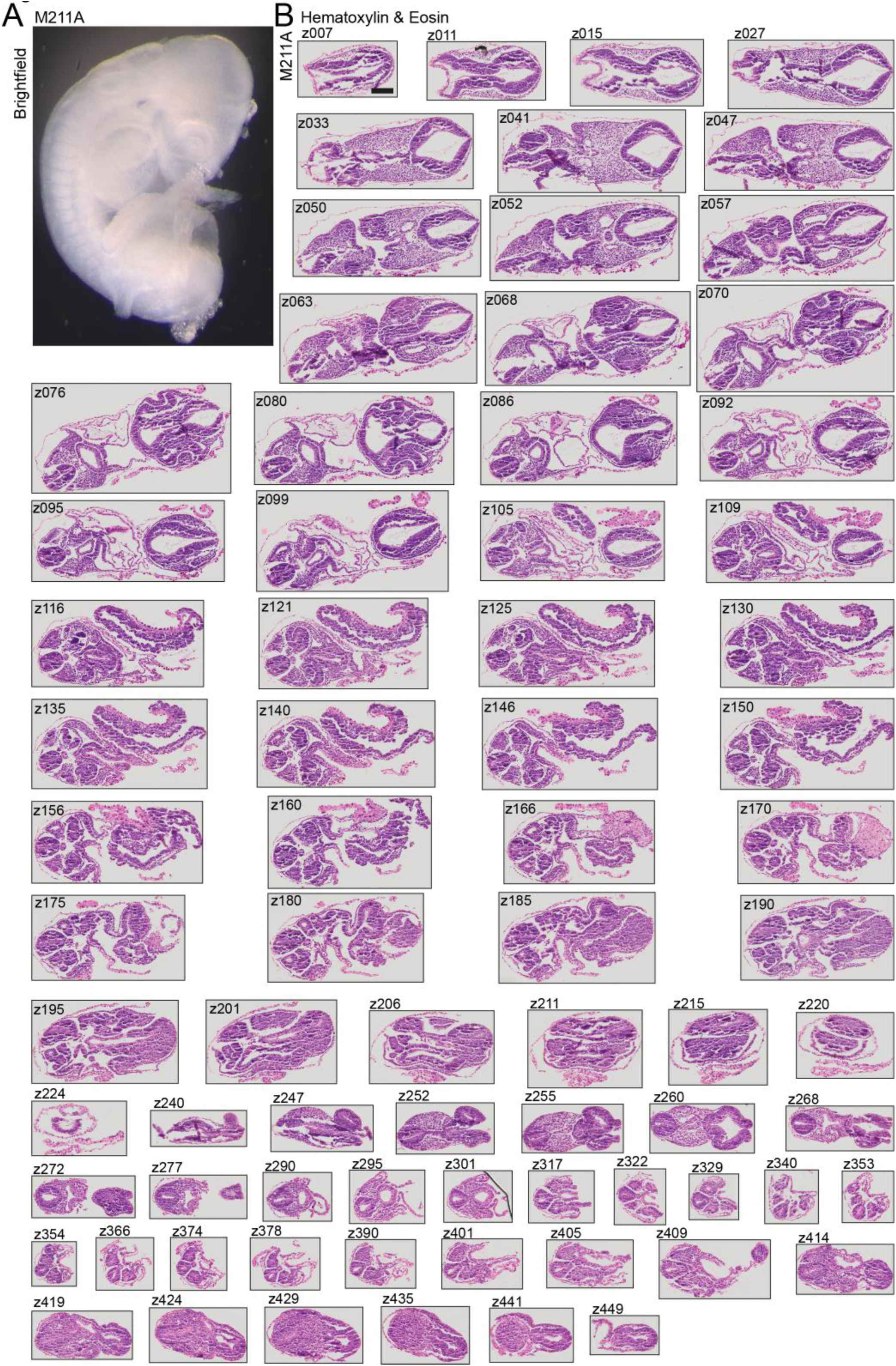
Cross sections of organogenesis embryo M211A. **A.** Brightfield images of right and left side of M211A. no scale bar. **B.** Hematoxylin and Eosin staining of paraffin cross sections of M211A. Cross section number is annotated in figure (z007-z449). Scale bars 100um. All imaged have the same scale bar.

**Figure S14:**
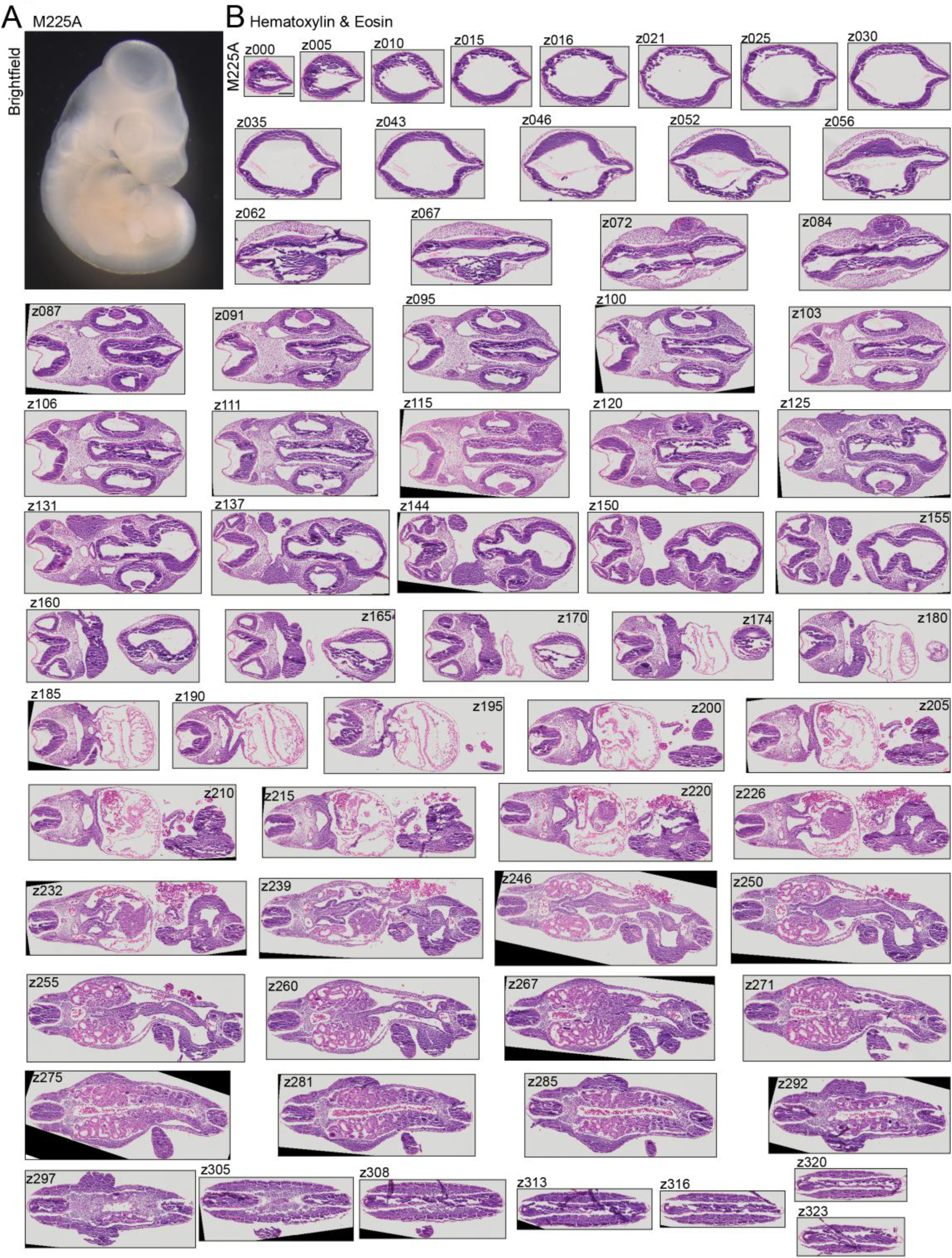
Cross sections of organogenesis embryo 225A. **A.** Brightfield images of right and left side of M225A. no scale bar. **B.** Hematoxylin and Eosin staining of paraffin cross sections of M225A. Cross section number is annotated in figure (z000-z323). Scale bar: 100um. All imaged have the same scale bar.

## Supplementary Tables

**Supplementary Table 1:** Staging Table of Pre-oviposition Development.

## Notes

### Competing Interest Statement

The authors have declared no competing interest.

## References

(1) White, M. E. Oogenesis and Early Embryogenesis. In Reproductive Biology and Phylogeny of Snakes; CRC Press, 2011.

(2) Eyal-Giladi, H.; Kochav, S. From Cleavage to Primitive Streak Formation: A Complementary Normal Table and a New Look at the First Stages of the Development of the Chick: I. General Morphology. Dev. Biol. 1976, 49 (2), 321–337. 10.1016/0012-1606(76)90178-0.

(3) Hamburger, V.; Hamilton, H. L. A Series of Normal Stages in the Development of the Chick Embryo. Dev. Dyn. 1992, 195 (4), 231–272. 10.1002/aja.1001950404.

(4) Wise, P. A. D.; Vickaryous, M. K.; Russell, A. P. An Embryonic Staging Table for In Ovo Development of Eublepharis Macularius, the Leopard Gecko. Anat. Rec. 2009, 292 (8), 1198–1212. 10.1002/ar.20945.

(5) Dufaure, J. P.; Hubert, J. Table de Developpement Du Lezard Vivipare: Lacerta Zootoca Vivipara Jacquin. Arch Anat Microsc Morphol Exp 1961, No. 50, 309–328.

(6) El Mouden, E.; Bons, J.; Pieau, C.; Renous, S.; Znari, M.; Boumezzough, A. Table de Développement Embryonnaire d’un Lézard Agamidé, Agama Impalearis Boettger, 1874. Ann. Sci. Nat. – Zool. Biol. Anim. 2000, 21 (3), 93–115. 10.1016/S0003-4339(00)01021-2.

(7) Blanc, F. Table de Développement de Chamaeleo Lateralis Gray, 1831. Ann Embryol Morphol 1974, No. 7, 99–115.

(8) Peter, K. Normentafel Zur Entwicklungsgeschichte Der Zauneidechse, Lacerta Agilis; Normentafeln zur Entwicklungsgeschichte der Wirbelthiere; Fischer: Berlin, 1904; Vol. 4.

(9) Levine, B. A.; Schuett, G. W.; Booth, W. Exceptional Long-Term Sperm Storage by a Female Vertebrate. PLOS ONE 2021, 16 (6), e0252049. 10.1371/journal.pone.0252049.

(10) Uller, T.; Olsson, M. Multiple Paternity in Reptiles: Patterns and Processes. Mol. Ecol. 2008, 17 (11), 2566–2580. 10.1111/j.1365-294X.2008.03772.x.

(11) Eckalbar, W. L.; Lasku, E.; Infante, C. R.; Elsey, R. M.; Markov, G. J.; Allen, A. N.; Corneveaux, J. J.; Losos, J. B.; DeNardo, D. F.; Huentelman, M. J.; Wilson-Rawls, J.; Rawls, A.; Kusumi, K. Somitogenesis in the Anole Lizard and Alligator Reveals Evolutionary Convergence and Divergence in the Amniote Segmentation Clock. Dev. Biol. 2012, 363 (1), 308–319. 10.1016/j.ydbio.2011.11.021.

(12) Tollis, M.; Hutchins, E. D.; Kusumi, K. Reptile Genomes Open the Frontier for Comparative Analysis of Amniote Development and Regeneration. Int. J. Dev. Biol. 2015, 58 (10-11–12), 863–871. 10.1387/ijdb.140316kk.

(13) Gilbert, S. F. Early Development in Birds. In Developmental Biology*. 6th edition*; Sinauer Associates, 2000.

(14) Sheng, G. Day-1 Chick Development. Dev. Dyn. 2014, 243 (3), 357–367. 10.1002/dvdy.24087.

(15) Bachvarova, R. F.; Skromne, I.; Stern, C. D. Induction of Primitive Streak and Hensen’s Node by the Posterior Marginal Zone in the Early Chick Embryo. Development 1998, 125 (17), 3521–3534. 10.1242/dev.125.17.3521.

(16) Cui, C.; Yang, X.; Chuai, M.; Glazier, J. A.; Weijer, C. J. Analysis of Tissue Flow Patterns during Primitive Streak Formation in the Chick Embryo. Dev. Biol. 2005, 284 (1), 37–47. 10.1016/j.ydbio.2005.04.021.

(17) Losos, J. B. Meet the Anoles! In Lizards in an Evolutionary Tree: Ecology and Adaptive Radiation of Anoles; Losos, J., Ed.; University of California Press, 2009; p 0. 10.1525/california/9780520255913.003.0002.

(18) Sanger, T. J.; Kircher, B. K. Model Clades Versus Model Species: Anolis Lizards as an Integrative Model of Anatomical Evolution. In Avian and Reptilian Developmental Biology: Methods and Protocols; Sheng, G., Ed.; Springer: New York, NY, 2017; pp 285–297. 10.1007/978-1-4939-7216-6_19.

(19) Geneva, A. J.; Park, S.; Bock, D. G.; de Mello, P. L. H.; Sarigol, F.; Tollis, M.; Donihue, C. M.; Reynolds, R. G.; Feiner, N.; Rasys, A. M.; Lauderdale, J. D.; Minchey, S. G.; Alcala, A. J.; Infante, C. R.; Kolbe, J. J.; Schluter, D.; Menke, D. B.; Losos, J. B. Chromosome-Scale Genome Assembly of the Brown Anole (Anolis Sagrei), an Emerging Model Species. *Commun*. Biol. 2022, 5 (1), 1–13. 10.1038/s42003-022-04074-5.

(20) Alföldi, J.; Di Palma, F.; Grabherr, M.; Williams, C.; Kong, L.; Mauceli, E.; Russell, P.; Lowe, C. B.; Glor, R. E.; Jaffe, J. D.; Ray, D. A.; Boissinot, S.; Shedlock, A. M.; Botka, C.; Castoe, T. A.; Colbourne, J. K.; Fujita, M. K.; Moreno, R. G.; ten Hallers, B. F.; Haussler, D.; Heger, A.; Heiman, D.; Janes, D. E.; Johnson, J.; de Jong, P. J.; Koriabine, M. Y.; Lara, M.; Novick, P. A.; Organ, C. L.; Peach, S. E.; Poe, S.; Pollock, D. D.; de Queiroz, K.; Sanger, T.; Searle, S.; Smith, J. D.; Smith, Z.; Swofford, R.; Turner-Maier, J.; Wade, J.; Young, S.; Zadissa, A.; Edwards, S. V.; Glenn, T. C.; Schneider, C. J.; Losos, J. B.; Lander, E. S.; Breen, M.; Ponting, C. P.; Lindblad-Toh, K. The Genome of the Green Anole Lizard and a Comparative Analysis with Birds and Mammals. Nature 2011, 477 (7366), 587–591. 10.1038/nature10390.

(21) Tollis, M.; Hutchins, E. D.; Stapley, J.; Rupp, S. M.; Eckalbar, W. L.; Maayan, I.; Lasku, E.; Infante, C. R.; Dennis, S. R.; Robertson, J. A.; May, C. M.; Crusoe, M. R.; Bermingham, E.; DeNardo, D. F.; Hsieh, S.-T. T.; Kulathinal, R. J.; McMillan, W. O.; Menke, D. B.; Pratt, S. C.; Rawls, J. A.; Sanjur, O.; Wilson-Rawls, J.; Wilson Sayres, M. A.; Fisher, R. E.; Kusumi, K. Comparative Genomics Reveals Accelerated Evolution in Conserved Pathways during the Diversification of Anole Lizards. Genome Biol. Evol. 2018, 10 (2), 489–506. 10.1093/gbe/evy013.

(22) Pirani, R. M.; Arias, C. F.; Charles, K.; Chung, A. K.; Curlis, J. D.; Nicholson, D. J.; Vargas, M.; Cox, C. L.; McMillan, W. O.; Logan, M. L. A High-Quality Genome for the Slender Anole (Anolis Apletophallus): An Emerging Model for Field Studies of Tropical Ecology and Evolution. G3 GenesGenomesGenetics 2024, 14 (1), jkad248. 10.1093/g3journal/jkad248.

(23) Kanamori, S.; Díaz, L. M.; Cádiz, A.; Yamaguchi, K.; Shigenobu, S.; Kawata, M. Draft Genome of Six Cuban Anolis Lizards and Insights into Genetic Changes during Their Diversification. BMC Ecol. Evol. 2022, 22 (1), 129. 10.1186/s12862-022-02086-7.

(24) Sanger, T. J.; Losos, J. B.; Gibson-Brown, J. J. A Developmental Staging Series for the Lizard Genus Anolis: A New System for the Integration of Evolution, Development, and Ecology. J. Morphol. 2008, 269 (2), 129–137. 10.1002/jmor.10563.

(25) Andrews, R. M. Oviposition Frequency of Anolis Carolinensis. Copeia 1985, 1985 (1), 259–262. 10.2307/1444828.

(26) Andrews, R.; Rand, A. S. Reproductive Effort in Anoline Lizards. Ecology 1974, 55 (6), 1317–1327. 10.2307/1935459.

(27) Lee, J. C.; Clayton, D.; Eisenstein, S.; Perez, I. The Reproductive Cycle of Anolis Sagrei in Southern Florida. Copeia 1989, 1989 (4), 930–937. 10.2307/1445979.

(28) Sever, D. M.; Hamlett, W. C. Female Sperm Storage in Reptiles. J. Exp. Zool. 2002, 292 (2), 187–199. 10.1002/jez.1154.

(29) Kircher, B. K.; Stanley, E. L.; Behringer, R. R. Anatomy of the Female Reproductive Tract Organs of the Brown Anole (Anolis Sagrei). Anat. Rec. 2024, 307 (2), 395–413. 10.1002/ar.25293.

(30) Kolbe, J. J.; Glor, R. E.; Rodríguez Schettino, L.; Lara, A. C.; Larson, A.; Losos, J. B. Genetic Variation Increases during Biological Invasion by a Cuban Lizard. Nature 2004, 431 (7005), 177–181. 10.1038/nature02807.

(31) Campbell, T. Northern Range Expansion of the Brown Anole (Anolis Sagrei) in Florida and Georgia. Herpetological review 1996, 27 (3), 155–157.

(32) Guraya, S. S. Ovarian Follicles in Reptiles and Birds; Zoophysiology; Springer: Berlin Heidelberg, 1989; Vol. 24.

(33) Vieira, S.; de Pérez, G. R.; Ramírez-Pinilla, M. P. Ultrastructure of the Ovarian Follicles in the Placentotrophic Andean Lizard of the Genus Mabuya (Squamata: Scincidae). J. Morphol. 2010, 271 (6), 738–749. 10.1002/jmor.10830.

(34) Silva, D. da; Cassel, M.; Mehanna, M.; Ferreira, A.; Dolder, M. A. H. Follicular Development and Reproductive Characteristics in Four Species of Brazilian Tropidurus Lizards. Zoolog. Sci. 2018, 35 (6), 553–563. 10.2108/zs180030.

(35) Jones, S. M. Chapter 4 – Hormonal Regulation of Ovarian Function in Reptiles. In Hormones and Reproduction of Vertebrates; Norris, D. O., Lopez, K. H., Eds.; Academic Press: London, 2011; pp 89–115. 10.1016/B978-0-12-374930-7.10004-4.

(36) Maurizii, M. G.; Taddei, C. Immunolocalization of Cytoskeletal Proteins in the Previtellogenic Ovarian Follicle of the Lizard Podarcis Sicula. Cell Tissue Res. 1996, 284 (3), 489–493. 10.1007/s004410050610.

(37) Magoffin, D. A. Ovarian Theca Cell. Int. J. Biochem. Cell Biol. 2005, 37 (7), 1344–1349. 10.1016/j.biocel.2005.01.016.

(38) Sandell, L. L.; Kurosaka, H.; Trainor, P. A. Whole Mount Nuclear Fluorescent Imaging: Convenient Documentation of Embryo Morphology. genesis 2012, 50 (11), 844–850. 10.1002/dvg.22344.

(39) Sandell, L.; Inman, K.; Trainor, P. DAPI Staining of Whole-Mount Mouse Embryos or Fetal Organs. Cold Spring Harb. Protoc. 2018, 2018 (10), pdb.prot094029. 10.1101/pdb.prot094029.

(40) Diaz Jr, R. E.; Shylo, N. A.; Roellig, D.; Bronner, M.; Trainor, P. A. Filling in the Phylogenetic Gaps: Induction, Migration, and Differentiation of Neural Crest Cells in a Squamate Reptile, the Veiled Chameleon (Chamaeleo Calyptratus). Dev. Dyn. 2019, 248 (8), 709–727. 10.1002/dvdy.38.

(41) E Gilland; Burke, A. Gastrulation in Reptiles. In Gastrulation: From cells to embryo.; Cold Spring Harbour Laboratory Press, 2004; pp 205–217.

(42) Trainor, P. A.; Tam, P. P. L. Cranial Paraxial Mesoderm and Neural Crest Cells of the Mouse Embryo: Co-Distribution in the Craniofacial Mesenchyme but Distinct Segregation in Branchial Arches. Development 1995, 121 (8), 2569–2582. 10.1242/dev.121.8.2569.

(43) Trainor, P. A.; Tan, S.-S.; Tam, P. P. L. Cranial Paraxial Mesoderm: Regionalisation of Cell Fate and Impact on Craniofacial Development in Mouse Embryos. Development 1994, 120 (9), 2397–2408. 10.1242/dev.120.9.2397.

(44) Noden, D. M.; Trainor, P. A. Relations and Interactions between Cranial Mesoderm and Neural Crest Populations. J. Anat. 2005, 207 (5), 575–601. 10.1111/j.1469-7580.2005.00473.x.

(45) Cuellar, O. Reproduction and the Mechanism of Meiotic Restitution in the Parthenogenetic Lizard Cnemidophorus Uniparens. J. Morphol. 1971, 133 (2), 139–165. 10.1002/jmor.1051330203.

(46) Smaga, C. R.; Bock, S. L.; Johnson, J. M.; Parrott, B. B. Sex Determination and Ovarian Development in Reptiles and Amphibians: From Genetic Pathways to Environmental Influences. Sex. Dev. 2022, 17 (2–3), 99–119. 10.1159/000526009.

(47) Tones, R. E.; Austin, H. B.; Lopez, K. H.; Rand, M. S.; Summers, C. H. Gonadotropin-Induced Ovulation in a Reptile (*Anolis Carolinensis*): Histological Observations. Gen. Comp. Endocrinol. 1988, 72 (2), 312–322. 10.1016/0016-6480(88)90214-6.

(48) Sheng, G. Epiblast Morphogenesis before Gastrulation. Dev. Biol. 2015, 401 (1), 17–24. 10.1016/j.ydbio.2014.10.003.

(49) Gilbert, S. F. Formation of the Neural Tube. In Developmental Biology*. 6th edition*; Sinauer Associates, 2000.

(50) Nikolopoulou, E.; Galea, G. L.; Rolo, A.; Greene, N. D. E.; Copp, A. J. Neural Tube Closure: Cellular, Molecular and Biomechanical Mechanisms. Development 2017, 144 (4), 552–566. 10.1242/dev.145904.

(51) Irie, N.; Kuratani, S. The Developmental Hourglass Model: A Predictor of the Basic Body Plan? Development 2014, 141 (24), 4649–4655. 10.1242/dev.107318.

(52) Pranter, R.; Feiner, N. Spatiotemporal Profiling of Neural Crest Cells in the Common Wall Lizard Podarcis Muralis. bioRxiv May 26, 2024, p 2024.05.24.595691. 10.1101/2024.05.24.595691.

(53) Hill, M. A. Embryology Carnegie Stages. https://embryology.med.unsw.edu.au/embryology/index.php/Carnegie_Stages (accessed 2024-07-11).

(54) Kaufmann, M. H. The Atlas of Mouse Development; 1992.

