## Supplementary Table 1 for "Pre-oviposition development of the brown anole (*Anolis sagrei*)"

| Stage | Overall process | Detail |
|---|---|---|
| 1 | | Zygote |
| 2 | Cleavage | first cleavage furrows, can be in various patterns |
| 3 | | cleavage divisions, group of cells - large in middle, furrows towards the outside of embryonic plate like spider web, cannot be recovered without plate of yolk beneath, easy to break apart |
| 4 | | small cells in middle, bigger towards the outside, still furrows extending to end of embryonic shield, cannot be recovered without plate of yolk beneath, easy to break apart |
| 5 | | tiny, bead-like cells in middle, getting a bit bigger towards outside, some furrows still extending, cells also into yolk, cannot be recovered without plate of yolk beneath, easy to break apart |
| 6 | perigastrulation | skin-like, flat, embryo can be peeled off like skin layer from yolk, very thin, flat. |
| 7 | | cosine shaping starts |
| 8 | | cosine pronounced, symmetrical |
| 9 | | midline emerges, left-right asymmetry |
| 10 | | midline full, left right asymmetry |
| 11 | Initiation of Neurulation | neural groove, completely open, folds elevated in middle but not too close, flat at anterior/posterior |
| 12 | | completely open, folds elevated and towards each other in middle, flat and open anterior/posterior |
| 13 | | middle close to each other or closed, anterior/posterior open, head folds flat open up to turned in but flat |
| 14 | | closed in middle, heat tiled, head folds start to bend towards each other, embryo straight, tail bud in angle |
| | | closed in middle, open anterior/posterior, stlightly bent all over, tail bud present |
| 15 | | closed in middle, anterior/posterior open, overall curled, head tilted, tail bud not curled, optic vesicles as slight swellings, heart starts |
| 16 | | anterior neuropore open, posterior closed, optic vesicles as swellings |
| | | anterior neuropore open, optic vesicles as swellings, small heart, future optic vesicle through HNK1 |

| | | |
|---|---|---|
| 17 | Completion of Neurulation | anterior neuropore open, optic vesicles as pronounced swellings, otic vesicle starting, heart as small sac, embryo straight, head tilted, initiation of pharyngal arch 1 |
| 18 | | anterior neuropore open, optic vesicles as swellings, otic vesicle present, 1 pharyngal arch, heart small, looped, body starts overall curling |
| 19 | | closed, optic vesicles pronounced, otic vesicle present, olfactory placode initiates, 1 pharyngal arch, heart small and looped |
| 20 | | closed, no invagination up to lens developing with wide optic fissure, otic vesicle, olfactory placode, 1-2 pharyngal arches, looped and large heart, overall curled, tail bud not curling yet or starting to be tilted but no tail yet |
| 21 | | closed, optic fissure open, otic vesicle, olfactory vesicle, 2-3 pharyngal arches, heard large for body, tail starts curling from bud to tail, kidney starts being visible |
|  |  | closed, optic fissure open, otic vesicle, olfactory placode, 2-3 pharyngal arches, overall curled, tail starts curling, kidney visible, heart looped, large for body |
| 22 | Organogenesis | closed, optic fissure open, otic vesicle, olfactory vesicle present, 3 pharyngal arches, heart looped and large, overall bent, tail curling, kidney visible |
| 23 | | closed, initiation limb buds, optic fissure open, otic vesicle/olfactory placode, 3 pharyngal arches, heart looped, large, kidney visible, overall turned |
| | | closed, initiation limb buds, optic fisure open, otic vesicle/olfactory placode, 3 pharyngal arches, heart looped and large |
| 24 | | closed, limb buds initiation, optic fissure open, otic vesicle present, olfactory placode present, 3-4 pharyngal arches, heart looped and large, overall curled, kidney nice, tail curled |
| | | closed, limb initiation, optic fissure open, otic vesicle/olfactory placode present, 3-4 pharyngal arches, heart large and looped, tail curled, overall curled |
| | | closed, limb buds initiation, optic fissure almost closed, otic vesicle, olfactory placode, 3-4 pharyngal arches, heaert large and looped, tail curled, overall curled, kidney nice |

| | | |
|---|---|---|
| 25 | | closed, front limb/hindlimb present, optic fissure almost closed, otic vesicle/olfactory placode, 4 pharyngal arches, heart very large and looped, tail curled, overall curled, kidney very pronounced |
| 26 | | closed, limbs pronounced, optic fissure almost closed, otic vesicle, olfactory placide, 4-5 pharyngal arches, heart large and looped, all curled, tail curled, kidney clear |
| | | closed, pronounced limbs, optic fissure almost closed, otic vesicle, olfactory placode, 4-5 pharyngal arches, heart looped and large, kidney pronounced, overall curled, tail curled |

| Somites |
| --- |
| 1 |
| 2 |
| 3 |
| 4 |
| 5 |
| 6 |
| 7 |
| 8 |

| |
|---|
| 9 |
| 13 |
| 14 |
| 15 |
| 18 |
| 19 |
| 20 |
| 22 |
| 25 |
| 26 |
| 27 |
| 28 |

| |
|---|
| 31 |
| 37 |
| 38 |
